# The dietary emulsifier polysorbate-80 induces lipid accumulation and cell death in intestinal epithelial cells via ferroptosis

**DOI:** 10.1101/2025.08.29.673091

**Authors:** Gonzalo Saiz-Gonzalo, Raminder Singh, Naomi Hanrahan, Gaston Cluzel, Cian Manning, Karina Quilter, Tadhg Crowley, Dagmar Srutkova, Tomas Hudcovic, Martin Schwarzer, Simone Marcone, Jacintha O’Sullivan, Susan A Joyce, Silvia Melgar

**Affiliations:** APC Microbiome Ireland, University College Cork, National University of Ireland, Cork, Ireland; Department of Medicine, University College Cork, National University of Ireland, Cork, Ireland; School of Biochemistry and Cell Biology, University College Cork, National University of Ireland, Cork, Ireland; Centre for Research in Vascular Biology, University College Cork, Cork, Ireland; Laboratory of Gnotobiology, Institute of Microbiology of the Czech Academy of Sciences, Nový Hrádek, Czech Republic; Department of Surgery, Trinity St. James’s Cancer Institute, Trinity Translational Medicine Institute, Trinity College Dublin, St. James’s Hospital, Dublin 8, Ireland

**Keywords:** Ferroptosis, IBD, polysorbate 80, oxidative stress, lipidomics, PUFAs, processed food

## Abstract

Chronic inflammatory and metabolic diseases are major global health issues increasingly linked to dietary factors. Consumption of dietary emulsifiers like polysorbate-80 (p80), common in ultra-processed foods and pharmaceuticals, has raised concerns about gut health. RNA sequencing on intestinal epithelial cells (IECs) exposed to p80 revealed increased expression of ferroptosis-associated genes and disruption of lipid metabolism pathways further demonstrated by mitochondrial dysfunction, including altered membrane potential and architecture, and accumulation of reactive oxygen species, iron, lipid peroxidation, and lipid droplet formation. Lipidomic profiling identified significant alterations in triglyceride species and elevated pro-ferroptotic polyunsaturated fatty acids. These data indicate that p80 disrupts lipid homeostasis in IECs and triggers ferroptotic cell death, mechanisms potentially contributing to the increased incidence of chronic conditions like inflammatory bowel disease and metabolic syndrome. The study highlights critical implications for public health, emphasizing the need for reassessment of emulsifier safety standards while balancing needs with consumer safety.

## Introduction

Chronic inflammatory and metabolic diseases, such as inflammatory bowel disease (IBD), metabolic syndrome, and certain forms of cancer, represent significant and growing public health concerns, affecting millions of individuals worldwide (Kaplan, 2015). Despite extensive research, the exact etiology of many chronic inflammatory and metabolic conditions remains unclear. Over the past few decades, however, the role of dietary habits in modulating disease risk has become increasingly evident (Bancil et al., 2021; Levine et al., 2018; Chassaing et al., 2017). Diet is now recognized as a major environmental factor that influences gut microbiota composition, host metabolic homeostasis, and mucosal and systemic inflammatory responses (Narula et al., 2021; Monteiro et al., 2019).

Westernised diet has been implicated in increasing the risk of these chronic conditions due to its high content of fats, sugars, read meat, processed food and food additives and low levels of fibre (Rizzello et al., 2019; Narula et al., 2021). One of the critical dietary components that has garnered attention is the widespread use of dietary emulsifiers, such as polysorbate-80 (p80). Emulsifiers are commonly used to stabilize the consistency of processed foods, such as cakes, ice creams, and chocolates. Interestingly, p80 alone is utilized in over 6000 pharmaceutical formulations, including treatments for Crohn’s disease (Nguyen et al., 2016; a et al., 2018). The inclusion of such additives has increased substantially over the past decades, paralleling the rise in processed and ultra-processed food consumption (Monteiro et al., 2019, Baker et al., 2020). Epidemiological studies have established correlations between the elements of the Westernized diet, including emulsifiers, and the heightened risk of developing IBD (Hou et al., 2011; Chapman-Kidell et al., 2010).

While p80 and other emulsifiers have been approved for use at concentrations up to 1% due to potential toxicity (Food Safety Commission, 2007), recent studies have revealed their deleterious effects on gut health. For example, dietary p80 and carboxymethylcellulose (CMC) induced low-grade inflammation, metabolic syndrome, and colitis in susceptible mice by disrupting the gut microbiota (notably depleting beneficial *Akkermansia* bacteria with anti-inflammatory properties) (Chassaing et al., 2015). Furthermore, p80 has been linked to the aggravation of conditions such as radiation enteritis in abdominal cancer patients (Li et al., 2020a). Despite these findings, the mechanism(s) by which p80 affects the intestine, including epithelial cells remain poorly understood.

Previous studies using rat tissues reported cell death induced by p80 (Tatsuishi et al., 2005; Yuan et al., 2015). This was corroborated by a previous study where p80 was shown to be toxic to intestinal epithelial cells (IECs) (Saiz-Gonzalo et al., 2021). Given the variety of mechanisms that regulate cell death, including necroptosis, pyroptosis, and ferroptosis (Galluzzi et al., 2015), the present study focused on identifying specific pathway(s) by which p80 induces IEC cell death. Among these, apoptosis is mediated by caspases and characterized by cell shrinkage, DNA fragmentation, and membrane blebbing (Elmore, 2007). Necroptosis is a caspase-independent necrotic cell death involving receptor-interacting protein kinases (RIPK1/RIPK3) and results in membrane rupture and release of damage-associated molecular patterns (Pasparakis & Vandenabeele, 2015). Pyroptosis involves inflammasome activation and gasdermin-mediated membrane pore formation, leading to inflammatory cell lysis (Man et al., 2017). In contrast, ferroptosis is characterised by iron-dependent oxidative stress, lipid peroxidation, and mitochondrial dysfunction, without the nuclear fragmentation typically associated with apoptosis (Dixon et al., 2012; Friedmann-Angeli et al., 2014).

In addition, an accumulation of lipid peroxides within polyunsaturated fatty acid (PUFA)-containing phospholipids are associated to ferroptotic cell death. Given the role of diet in modulating gut inflammation and the observed toxicity of p80, we hypothesized that p80 might induce cell death in IECs, contributing to the pathogenesis of several conditions including IBD. To explore this hypothesis, an RNASeq analysis was conducted followed by an in-depth investigation into the effects of p80 on lipid metabolism and cell death pathways in IECs. Findings from this study indicate p80 exposure resulted in an early mitochondria dysfunction, characterised by accumulation of reactive oxygen species (ROS), dysregulated electric mitochondria potential, accumulation of Fe, followed by increased lipid peroxidation resulting in accumulation of lipid droplets and subsequent cell death. Addition of small molecule inhibitors targeting ferroptosis revealed that p80 indeed induced ferroptotic cell death in IECs. Lipidomic analysis revealed significant alterations in triglycerides and pro-ferroptotic PUFAs, which were partly recovered by the addition of ferroptosis inhibitors. In conclusion, these findings suggest that p80-triggered IEC ferroptosis may compromise barrier integrity and promote metabolic endotoxemia, potentiating the low-grade inflammation and dysbiosis that underlie diet-induced obesity and IBD.

## RESULTS

### Polysorbate 80 induces cytotoxicity and membrane permeability alterations in epithelial cells and intestinal organoids

To investigate the effects of p80 on intestinal epithelial cell viability, HT29, C2Bbe1 and DLD1 cells were treated with various concentrations of p80 (1, 0.25%, 0.125%, 0.06%, 0.001%) for 6, and 10 hours (Supplementary Figure 1a). Epithelial cells exposed to the FDA-approved concentration of 1% p80 exhibited 90% cell death within 6 hours (U.S. FDA, 2023; Supplementary Figure 1a). Assessment using Sytox Green staining demonstrated a significant increase in membrane permeability, indicative of cell death (Veldhuis et al., 2001), in a time and dose-dependent manner (Figure 1a,b). At the second highest concentration (0.25%), a significant decrease in cell viability was observed as early as 3 hours, with pronounced effects noted by 10 hours, suggesting rapid cytotoxicity induced by p80.

**Figure 1.**
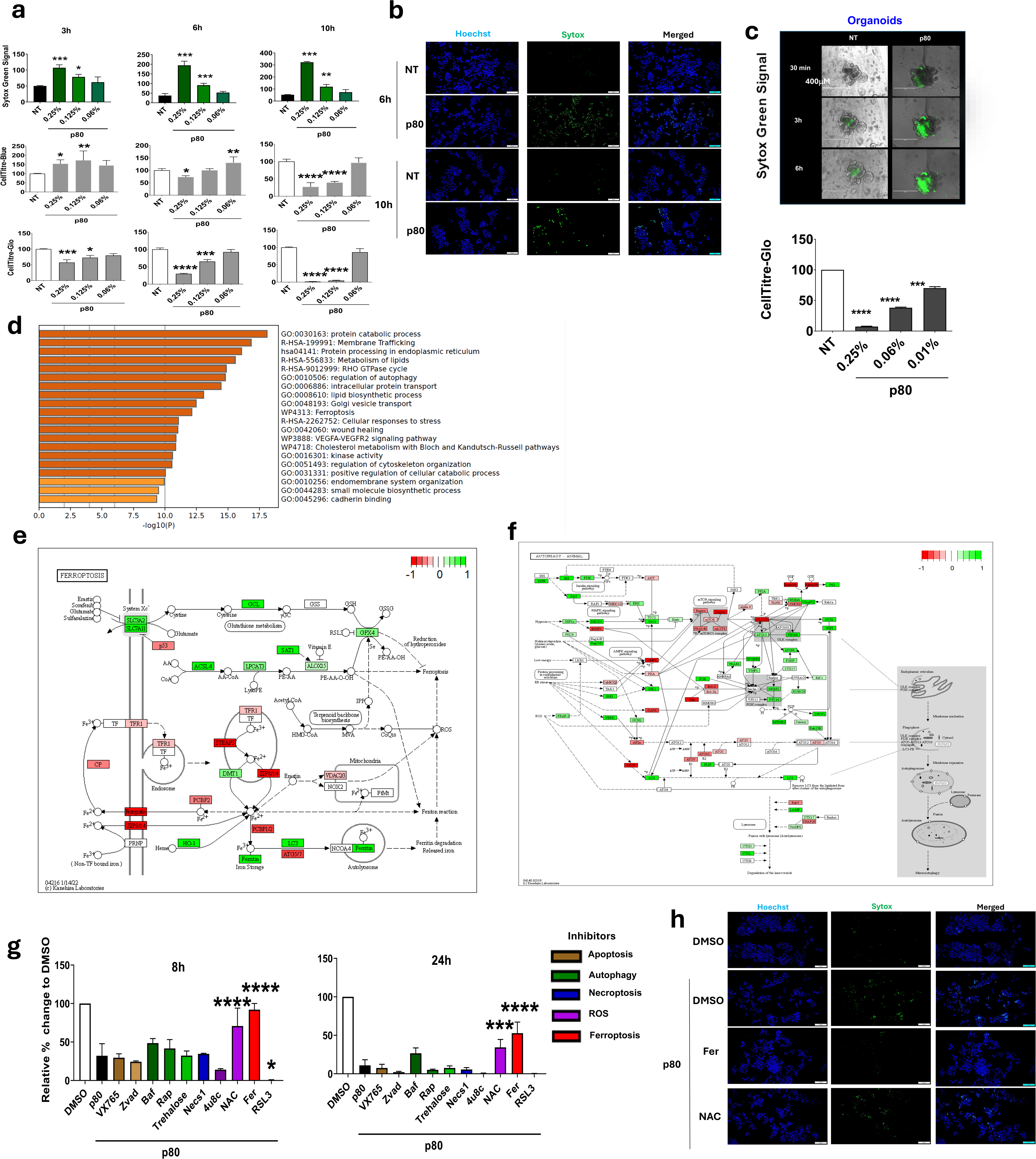
Polysorbate 80 induces cytotoxicity and cell death in intestinal epithelial cells and organoids. **(a)** Time- and dose-dependent loss of HT29 viability after treatment with p80 (0–0.25 %) for 3–10 h quantified by Sytox Green uptake, CellTiter-Blue and -Glo assays. **(b)** Sytox Green micrographs of HT29 cells exposed to 0.25 % p80 (6 h, 10 h). Scale bar, 400 µm. **(c)** Murine intestinal organoids treated with p80 (0.01–0.25 %) for 30 min–6 h: Sytox Green images (up; scale bar, 400 µm) and CellTiter-Glo viability (down). **(d)** GO and KEGG enrichment of differentially expressed genes in HT29 cells after 2 h 0.25% p80 treatment. **(e, f)** KEGG ferroptosis (e) and autophagy (f) pathways map highlighting p80-regulated genes. Nodes are colored by RNA-seq differential expression where red = upregulated transcripts (log₂FC > 0), green = downregulated transcripts (log₂FC < 0); grey = not detected/not significant (FDR < 0.05 where shown). **(g)** Rescue assay: viability of HT29 cells treated with p80 for 8 h and 24 h in the presence of apoptosis (VX-765, zVAD), autophagy (Rapamycin, Trehalose), necroptosis (Nec-1), ROS (NAC) or ferroptosis (Ferrostatin-1, RSL3) inhibitors. **(h)** Sytox Green staining of HT29 cells treated with 0.25% p80 for 8 h in the presence of Ferroptosis (Fer-1) and ROS (NAC) inhibitors and stained with Hoechst (blue) and Sytox Green (green). Scale bars: 400 μm. Data presented as mean ± SD. (n = 3). Statistical significance was assessed by one-way ANOVA with Tukey’s post-hoc test (*p < 0.05; **p < 0.01; ***p < 0.001; ****p < 0.0001 versus non-treated, NT in a and vs p80 in g). Images are representative of ≥ 3 independent experiments.

To further confirm the impact of p80 on cell viability, CellTiter-Blue and CellTiter-Glo assays were conducted (Figure 1a). These assays corroborated the Sytox Green findings, showing a marked reduction in viability at the higher concentration of 0.25% p80. Notably, the viability of HT29 cells was significantly compromised as early as 6 hours post-treatment. Similar results were obtained with two other epithelial cell lines, C2Bbe1 and DLD1 cells (Supplementary Fig 1a). To examine whether the toxic effect was a feature of all dietary emulsifiers, carboxymethylcellulose (CMC) was also examined. In contrast to p80 CMC caused no toxic effect on IECs, however, at higher concentrations it reduced metabolic viability at 1 h and 24 h (Supplementary Figure 1b).

These findings were extended to a 3D culture model using murine intestinal organoids to explore the translational relevance of p80-induced cytotoxicity (Figure 1c). Mature organoids treated with 0.01% p80 showed significant structural disintegration and increased membrane permeability over time, with the highest alteration at 3 and 6 hours, as evidenced by Sytox Green staining (Figure 1c) and supported by the reduced cell viability using CellTiter-Glo at the tested p80 concentrations at 6 hours (Figure 1c). In addition, the development of the organoids was also compromised during an 8-day time-lapse culture in p80, characterised by an earlier (day 6) disintegration of the organoid exposed to 0.06% p80 (Supplementary Figure 1c).

Further mechanistic insights were gained by bulk RNASeq on HT29 cells treated with 0.25% p80 for 2 hours. Gene ontology (GO) enrichment analysis revealed significant upregulation of pathways related to lipid metabolism, autophagy, ferroptosis, and the cellular stress response (Figure 1d). Using KEGG pathway mapping, we overlaid the p80-regulated DEGs onto the ferroptosis pathway and found significant perturbations in modules governing iron handling and glutathione/GPX4 redox metabolism (Figure 1e). Whereas autophagy-pathway mapping revealed ER/UPR activation together with down-regulation of mTOR-signaling components and several ATG transcripts (Figure 1f). To functionally test these pathways, HT29 cells were pre-treated with modulators of ferroptosis (Ferrostatin-1, RSL3), autophagy (rapamycin, trehalose), ROS (NAC), apoptosis (VX-765, zVAD-fmk), or necroptosis (Nec-1) prior to p80 exposure, and viability/membrane permeability were quantified relative to p80 alone (Figure 1g). In line with the non-recovery by the pan-apoptosis inhibitor zVAD, caspase-3/7 activity was not induced in cells treated with p80 for up to 24 hours (Supplementary Figure 2a). In contrast and in line with RNASeq findings, inhibitors such as Ferrostatin-1 (Fer) and N-acetylcysteine (NAC), which target ferroptosis and ROS pathways, respectively, significantly recovered cell viability at 8h (Figure 1g and h) and partially at 24h (Figure 1g).

### Polysorbate 80 induces ROS production, mitochondrial dysfunction, and lipid peroxidation in HT29 cells

Further analysis on the RNA-Seq data set identified significant upregulation of key genes involved in oxidative stress responses and cell death pathways, including the genes *ANXA1* (response to oxidative stress), *ERN1* (cellular response to reactive oxygen species) (Table 1). To further examine the influence of p80 on oxidative stress pathway and mitochondrial function in IECs, ROS production was assessed over a 24-hour period using flow cytometry (Figure 2a). A significant increase in ROS levels was observed in p80-treated cells compared to non-treated (NT) controls, with the highest ROS levels detected at 1-hours post-treatment followed by reduced levels over time.

**Figure 2.**
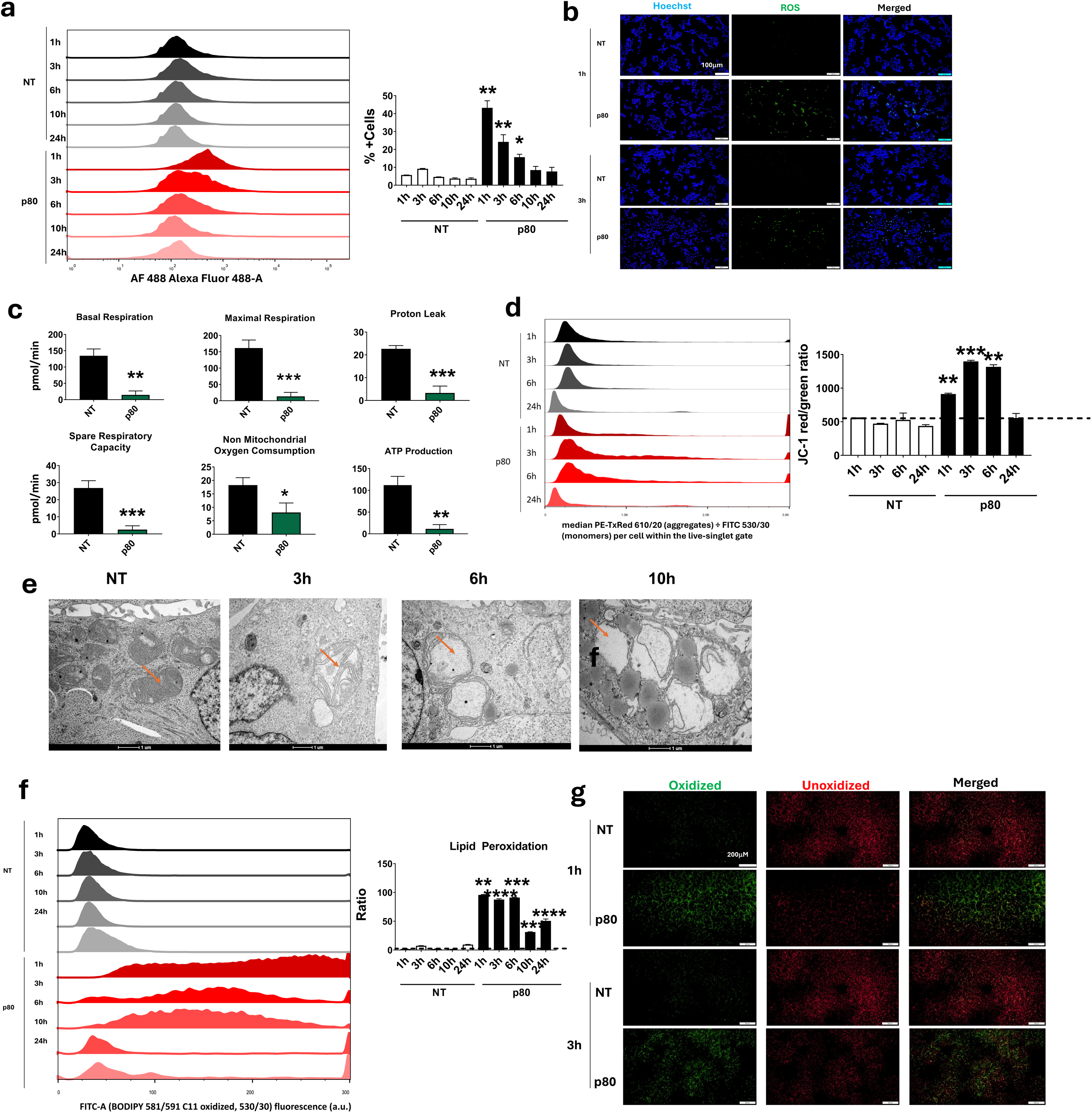
Polysorbate 80 triggers oxidative stress, mitochondrial dysfunction, and lipid peroxidation in epithelial cells. **(a)** ROS by H₂DCFDA flow cytometry. Left: FITC-A (H₂DCFDA, 488) fluorescence (a.u.) density histograms (bi-exponential) for live singlets after 0, 1, 3, 6, 10, 24 h with 0.25% p80 vs NT. Right: ROS MFI (FITC), normalized to NT (=1). % ROS-positive (FMO-defined) is provided in Supplementary Fig. 6. **(b)** Confocal ROS images of HT29 cells after 1 h and 3 h 0.25% p80; ROS, green; nuclei, Hoechst (blue). Scale bar, 100 µm. **(c)** Seahorse XF Cell Mito Stress Test after 3 h p80. Bars show OCR (pmol O₂·min⁻¹) parameters: basal respiration, maximal respiration, proton leak, spare respiratory capacity, non-mitochondrial OCR, and ATP production. **(d)** ΔΨm by JC-1 flow cytometry. Left: FITC (530/30; JC-1 monomers, green) and PE-TxRed (610/20; JC-1 aggregates, red) histograms (bi-exponential) for live singlets. Right: JC-1 red/green ratio = median(PE-TxRed 610/20 ÷ FITC 530/30) per cell, normalized to NT (=1). Detail on Supplementary Fig. 6. **(e)** Transmission electron microscopy (TEM) ultrastructure of HT29 cells treated with 0.25% p80 for 3, 6 and 10 h; orange arrows mark mitochondria structural changes. Scale bar, 1 µm. **(f)** Lipid peroxidation by BODIPY 581/591 C11. Left: density histograms (bi-exponential) for live singlets showing oxidized (FITC 530/30) and non-oxidized (PE 585/42) C11 signals over 0–24 h after 0.25% p80 vs NT). Detail on Supplementary Fig. 6. **(g)** Confocal BODIPY 581/591 C11 at 1, 3, and 6 h: oxidized (green, 510 nm) and non-oxidized (red, 590 nm); nuclei Hoechst. Scale bar, 200 µm. Unless stated otherwise, graphs show mean ± SD from ≥3 independent experiments. One-way ANOVA (Tukey): *p < 0.05, **p < 0.01, ***p < 0.001, ***p < 0.0001 vs NT.

**Table 1.** p80-induced key upregulated genes associated with oxidative stress pathways^a^.

| Gene | Description | LogP | GO Name | GO Number |
| --- | --- | --- | --- | --- |
| **AKR1C1** | aldo-keto reductase 1C1 | -33.00 | regulation of reactive oxygen species metabolic process | GO:2000377 |
| **ANXA1** | Annexin A1 | -46.72 | Response to oxidative stress | GO:0006979 |
| AREG | Amphiregulin | -46.63 | Response to reactive oxygen species | GO:0000302 |
| AXL | AXL receptor tyrosine kinase | -38.71 | Response to hydrogen peroxide | GO:0042542 |
| BAK1 | Bcl-2 homologous antagonist/killer | -35.04 | Chemical carcinogenesis - reactive oxygen species | hsa05208 |
| CAT | Catalase | -34.99 | Cellular response to oxidative stress | GO:0034599 |
| CYP1B1 | Cytochrome P450 family 1 subfamily B member 1 | -21.25 | Positive regulation of cell death | GO:0010942 |
| **EPHX1** | epoxide hydrolase 1 | -9.03 | Photodynamic therapy-induced NFE2L2 (NRF2) survival signaling | WP3612 |
| ERN1 | Endoplasmic reticulum to nucleus signaling 1 | -26.59 | Cellular response to reactive oxygen species | GO:0034614 |
| FOSL1 | Fos-related antigen 1 | -17.30 | DNA damage response (only ATM dependent) | WP710 |
| HIF1A | Hypoxia inducible factor 1 subunit alpha | -20.85 | Positive regulation of programmed cell death | GO:0043068 |
| JUN | Jun proto-oncogene, AP-1 transcription factor | -24.27 | Cellular response to hydrogen peroxide | GO:0070301 |
| **MAPK8** | Mitogen-activated protein kinase 8 | -18.70 | Colorectal cancer | hsa05210 |
| NQO1 | NAD(P)H quinone dehydrogenase 1 | -33.00 | Regulation of reactive oxygen species metabolic process | GO:2000377 |
| PLK3 | Polo-like kinase 3 | -34.72 | Cellular response to chemical stress | GO:0062197 |
| PPARGC1B | Peroxisome proliferator-activated receptor gamma coactivator 1-beta | -18.30 | Pathways in cancer | hsa05200 |
| RHOB | Ras homolog family member B | -39.18 | Response to inorganic substance | GO:0010035 |
| RIPK3 | Receptor interacting serine/threonine kinase 3 | -15.10 | Regulation of epithelial cell migration | GO:0010632 |
| SIRT1 | Sirtuin 1 | -19.66 | Positive regulation of apoptotic process | GO:0043065 |
| THBS1 | Thrombospondin 1 | -15.27 | Gastrin signaling pathway | WP4659 |
| VEGFA | Vascular endothelial growth factor A | -15.14 | Response to decreased oxygen levels | GO:0036293 |
| ZC3H12A | Zinc finger CCCH-type containing 12A | -16.52 | MET in type 1 papillary renal cell carcinoma | WP4205 |

<sup>a</sup> Data generated from RNA-seq analysis on GO-term on HT29 cells cultured for 2 hours with 0.25% p80

Confocal microscopy studies supported the flow cytometry data by showing the intracellular generation of ROS in p80-treated cells at 1 and 3 hours (Figure 2b).

Given the role of mitochondria in cellular energy production and ROS regulation, the effect of p80 was evaluated on mitochondrial function using the Seahorse Mito Stress Test (Figure 2c). p80 treatment led to a significant reduction in Basal Respiration, Maximal Respiration, Spare Respiratory Capacity, and ATP Production, alongside a decrease in Proton Leak and Non-Mitochondrial Oxygen Consumption. These results indicate that p80 severely impairs mitochondrial respiration and energy production, likely contributing to the observed increase in ROS and cellular stress. To further examine the effect of p80 on the mitochondrial membrane potential health status, a JC-1 dye was used. A significant increase in JC-1 ratio at 1-, 3- and 6-hour time points were identified by flow cytometry (Figure 2d).

To further examine mitochondrial integrity, a kinetic transmission electron microscopy (TEM) study was conducted on p80-treated cells (Figure 2e). TEM images revealed marked mitochondrial damage in p80-treated cells, as early as 3 hours post-treatment, including disrupted cristae and mitochondrial swelling, consistent with the observed functional impairments (Figure 2a-d). These structural abnormalities are indicative of mitochondrial distress and dysfunction induced by p80.

Lipid peroxidation, a downstream effect of ROS-activation and a key feature of ferroptosis was assessed by flow cytometry over 24 hours (Figure 2f). A highly significant increase in lipid peroxidation ratio was detected in p80-treated cells, particularly at the 3- and 6-hour time points, paralleling the findings on ROS levels and demonstrating significant oxidative damage to cellular lipids.

Finally, confocal microscopy was used to visualize oxidized and unoxidized lipids in p80-treated cells (Figure 2g). Increased levels of oxidized lipids were evident at 1- and 3-hours post-treatment, further validating the occurrence of lipid peroxidation in response to p80 over the NT controls.

### Polysorbate 80 induces ferroptosis in epithelial cells through and iron accumulation

To investigate ferroptosis further, mitochondrial and cellular labile iron levels were assessed using MitoFerroGreen and FerroOrange staining, respectively, over a 24-hour period via flow cytometry. MitoFerroGreen staining and quantification (Figure 3a) revealed a significant rise in mitochondrial iron accumulation in p80-treated cells up to 10 hours post-p80 treatment, with the highest peak at 3 hours. Notably, ANXA1 (logP - 46.72357) and CAT (logP −34.98751) encoding proteins crucial for the cellular response to oxidative stress (Table 1), together with elevated ROS levels aligns with prior studies linking oxidative stress and iron dysregulation to ferroptosis in other cell types (Yang et al., 2016; Gao et al., 2016). Similarly, FerroOrange staining and quantification (Figure 3b) demonstrated elevated cellular labile iron levels, aligning with ferroptosis induction, as reported in studies emphasizing the role of labile iron in driving lipid peroxidation and ferroptosis (Stockwell et al., 2017; Li et al., 2020b). These findings highlight the involvement of iron accumulation in p80-induced ferroptosis and support the evidence implicating iron overload as a key mediator of ferroptotic cell death (Friedmann Angeli et al., 2014; Dixon et al., 2012).

**Figure 3.**
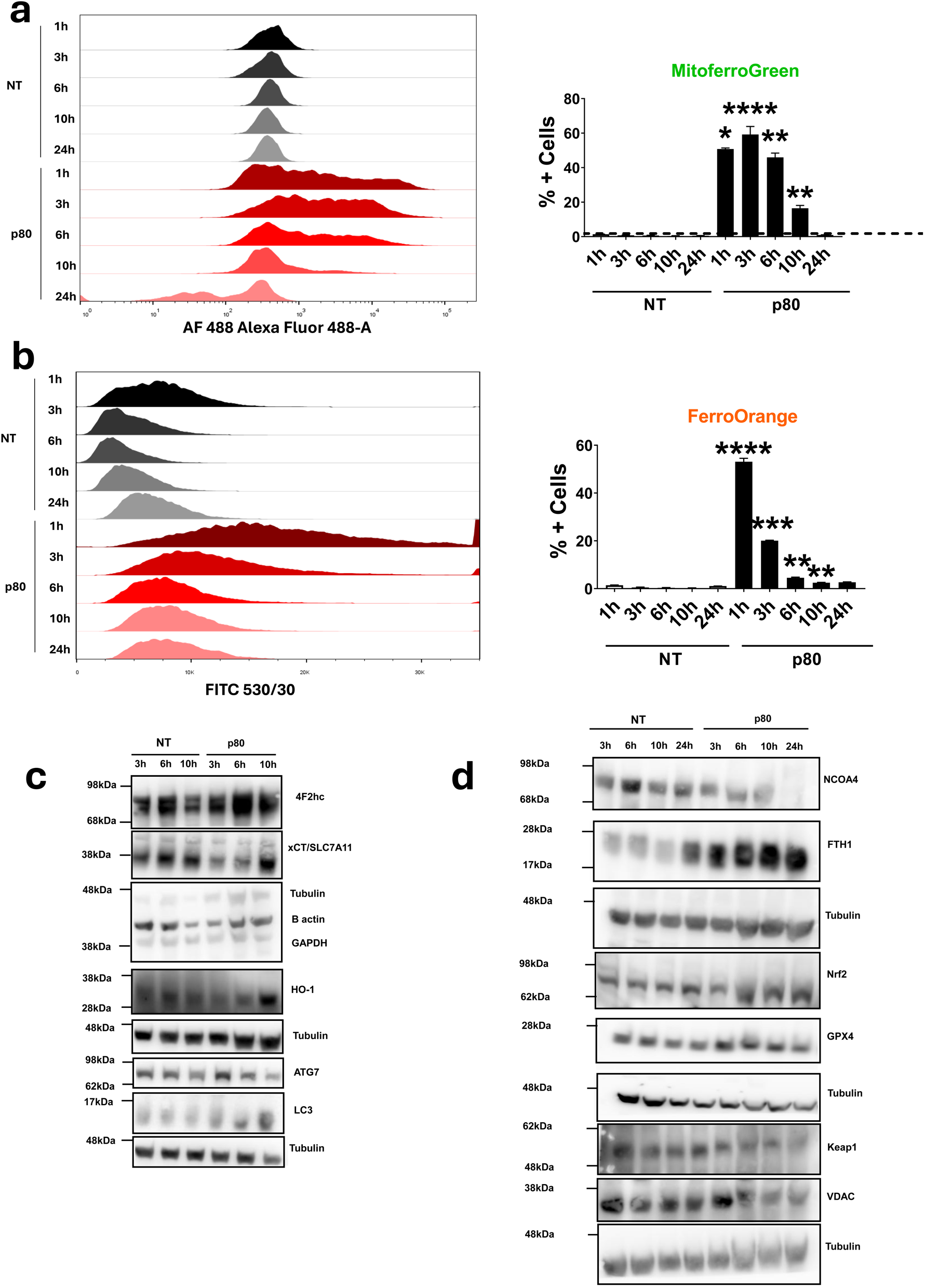
Polysorbate 80 (p80) induces ferroptosis through oxidative stress, mitochondrial dysfunction, and iron accumulation in HT29 cells. **(a)** Mitochondrial Fe²⁺ by Mito-FerroGreen (flow cytometry). Left: AF488-A (Mito-FerroGreen) fluorescence (a.u.) density histograms (bi-exponential) for live singlets at 1–24 h after 0.25% p80 vs NT. Right: % Mito-FerroGreen-positive (of live singlets); gate set with unstained + AF488 FMO; single-stain compensation applied. Percent data were analyzed after arcsine–square-root transform. Detailed provided in Supplementary Fig. 6. **(b)** Cytosolic labile iron levels measured by FerroOrange staining under 0.25 % p80 for 0–24 h; histograms (left) and quantification (percentage of positive cells, right) shown. Detailed provided in Supplementary Fig. 6. **(c, d)** Western blot analysis of ferroptosis-related proteins (4F2hc, xCT/SLC7A11, NCOA4, FTH1, GPX4) and oxidative stress markers (Nrf2, HO-1, Keap1) in HT29 cells treated with p80 over time; β-actin, GAPDH, and Tubulin are used as loading controls. Graphs show mean ± SD from three independent biological experiments. Statistical significance determined by one-way ANOVA with Tukey’s test (*p < 0.05; **p < 0.01; ***p < 0.001; ****p < 0.0001 versus non-treated, NT). Images are representative of ≥ 3 independent experiments.

Western blot and RNA-seq analysis confirmed the involvement of key ferroptosis and oxidative stress pathways in HT29 cells treated for X hrs with p80 (Figures 3c, d; Table 2 respectively). An upregulation of ferroptosis-related proteins, including 4F2hc and xCT/SLC7A11 (Figure 3c) and the iron-storage protein FTH1 (Figure 3d), was observed across 3, 6, 10, and 24 hours, consistent with RNA-seq results at the 6-hour time point (Table 2). Markers of oxidative stress such as HO-1 were similarly elevated (Figures 3c, d), while Nrf2 protein levels remained unchanged, despite increased NFE2L2 gene expression detected by RNA-seq (Table 2). HMOX1, a key enzyme in the heme catabolic process, was also upregulated at 6 hours (Table 2).

**Table 2.**
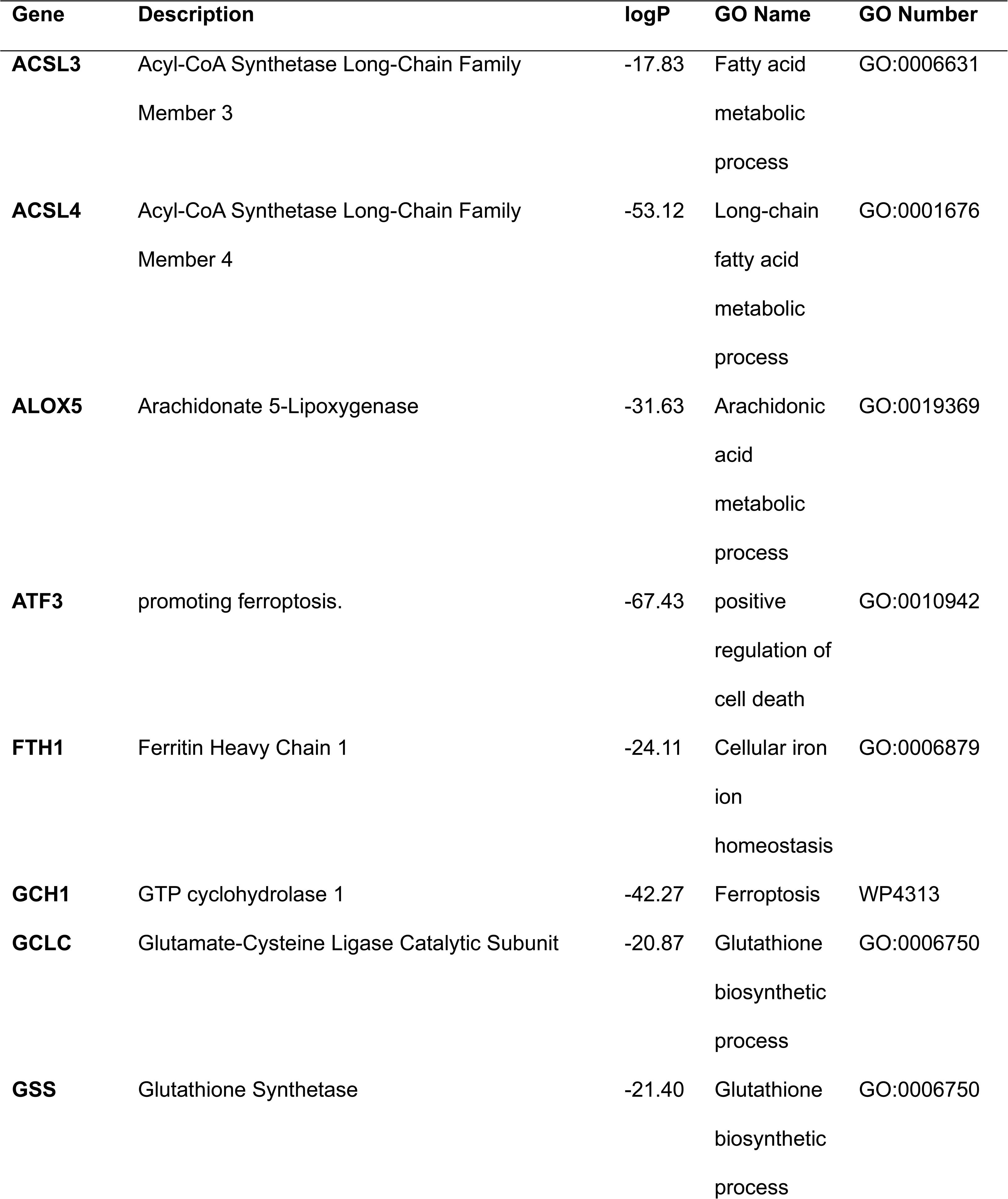
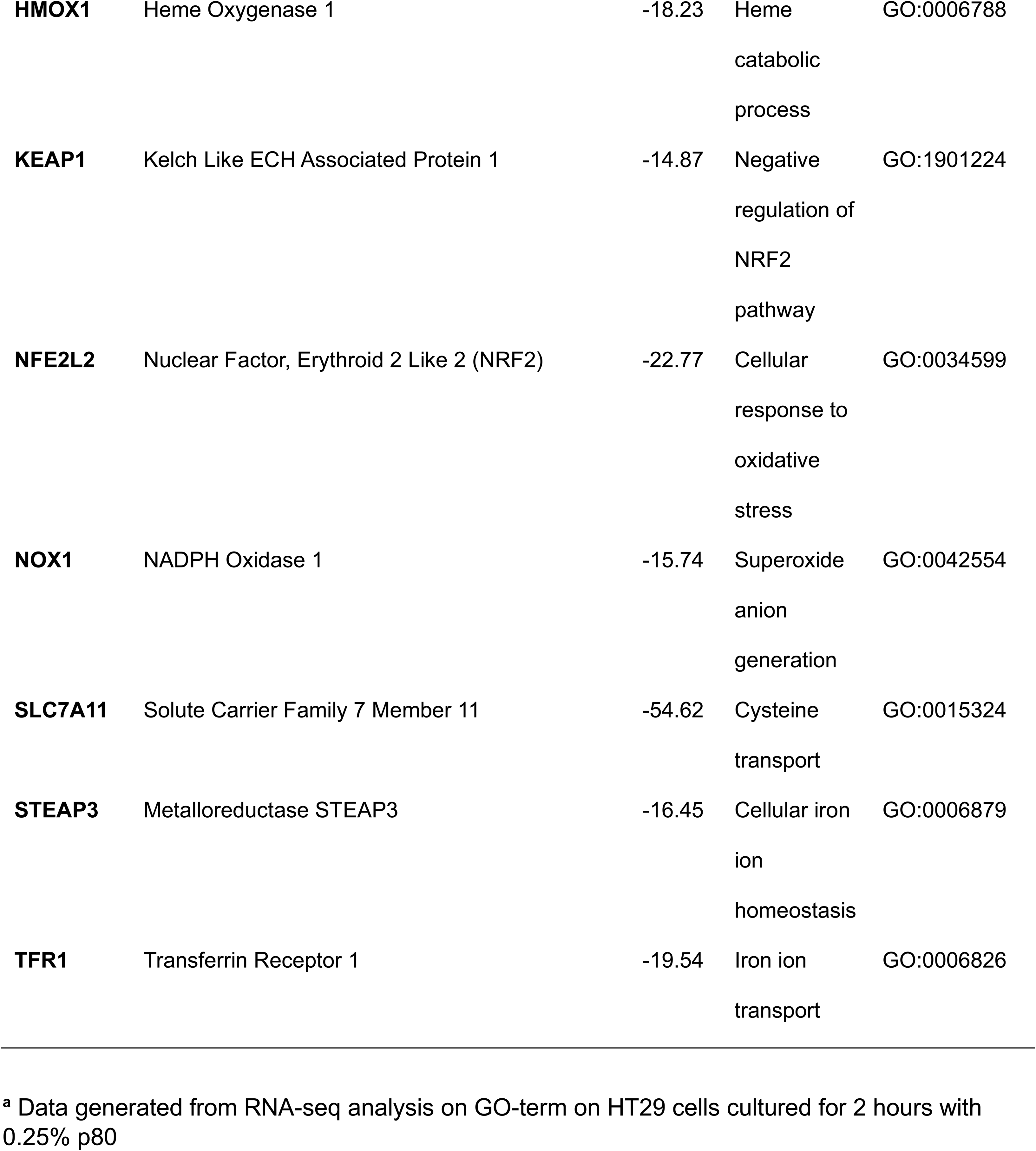
Analysis of RNA-seq data at the 6-hour time point of key upregulated genes involved in ferroptosis and oxidative stress pathways^a^.

At the earlier 2-hour time point, RNA-seq revealed increased expression of genes involved in oxidative stress response, including AREG and ERN1, along with the stress-related signaling molecules MAPK8 and FOSL1 (Table 1). Furthermore, upregulation of enzymes involved in lipid peroxidation metabolism, such as AKR1C1 and EPHX1, was observed at 6 hours (Table 2), indicating possible activation of antioxidant defenses as part of the ferroptotic response.

In contrast, a downregulation of NCOA4, Keap1, and VDAC proteins was detected over time (Figure 3), suggesting modulation of ferritinophagy, antioxidant regulation, and mitochondrial permeability (Mancias et al., 2014; Fan et al., 2017). A similar reduction of KEAP1 gene expression was observed at 6 hours (Figure 3d), consistent with decreased negative regulation of the NRF2 pathway (GO:1901224) (Table 2).

Notably, the expression of the canonical ferroptosis-associated gene GPX4 remained unchanged at both the RNA and protein levels (Figure 3d) (Friedmann-Angeli et al., 2014). Other key ferroptosis regulators assessed included ferroptosis suppressor protein 1 (FSP1) (Doll et al., 2019), dihydroorotate dehydrogenase (DHODH) (Mao et al., 2021), and MBOAT2 (Liang et al., 2023), showing no significant changes in the RNAseq analysis. In contrast, GTP cyclohydrolase 1 (GCH1), associated with the

GPX4-independent ferroptosis protection pathway (Kraft et al., 2020), was significantly upregulated (Table 2).

### Treatment with Ferrostatin-1 reduces ferroptosis in intestinal epithelial HT29 cells and murine intestinal loops

As shown earlier in Figure 1g, the ferroptosis inhibitor Ferrostatin-1 (Fer) recovered the p80-induced cell death. In addition to Ferrostatin-1 (Fer), Liproxstatin-1 (Lip), Tocopherol (Toco), Deferoxamine (DFO), and Zileuton (Zil), which target lipid peroxidation (Dixon et al., 2012), sulfasalazine (SAS) and rosiglitazone (Rosi), which inhibit ACSL4 (Chen et al., 2020), as well as the statins rosuvastatin and fluvastatin, which are reported to modulate oxidative stress and lipid metabolism (Haruna et al., 2007; Liu et al., 2020b) were examined in a kinetic study (Supplementary Figure 3).

At 3 hours post-treatment (Figure 4a), Fer significantly enhanced cell viability compared to p80 alone, aligning with our previous observations (Figure 1g). By 6 hours and 10 hours (Figure 4a), Fer, Lip, Toco, DFO, and Zil continued to confer a protective effect, improving cell viability in p80-treated cells and confirming that p80-induces cell death via ferroptosis. In contrast, sulfasalazine and the two statins (rosuvastatin, fluvastatin) reduced viability further than p80 alone, suggesting that under these conditions, they may exacerbate ferroptotic or oxidative damage (Supplementary Figure 3). Rosiglitazone also failed to restore viability, indicating that p80-induced ferroptotic cell death may not rely primarily via ACSL4 activation (Chen et al., 2020).

**Figure 4:**
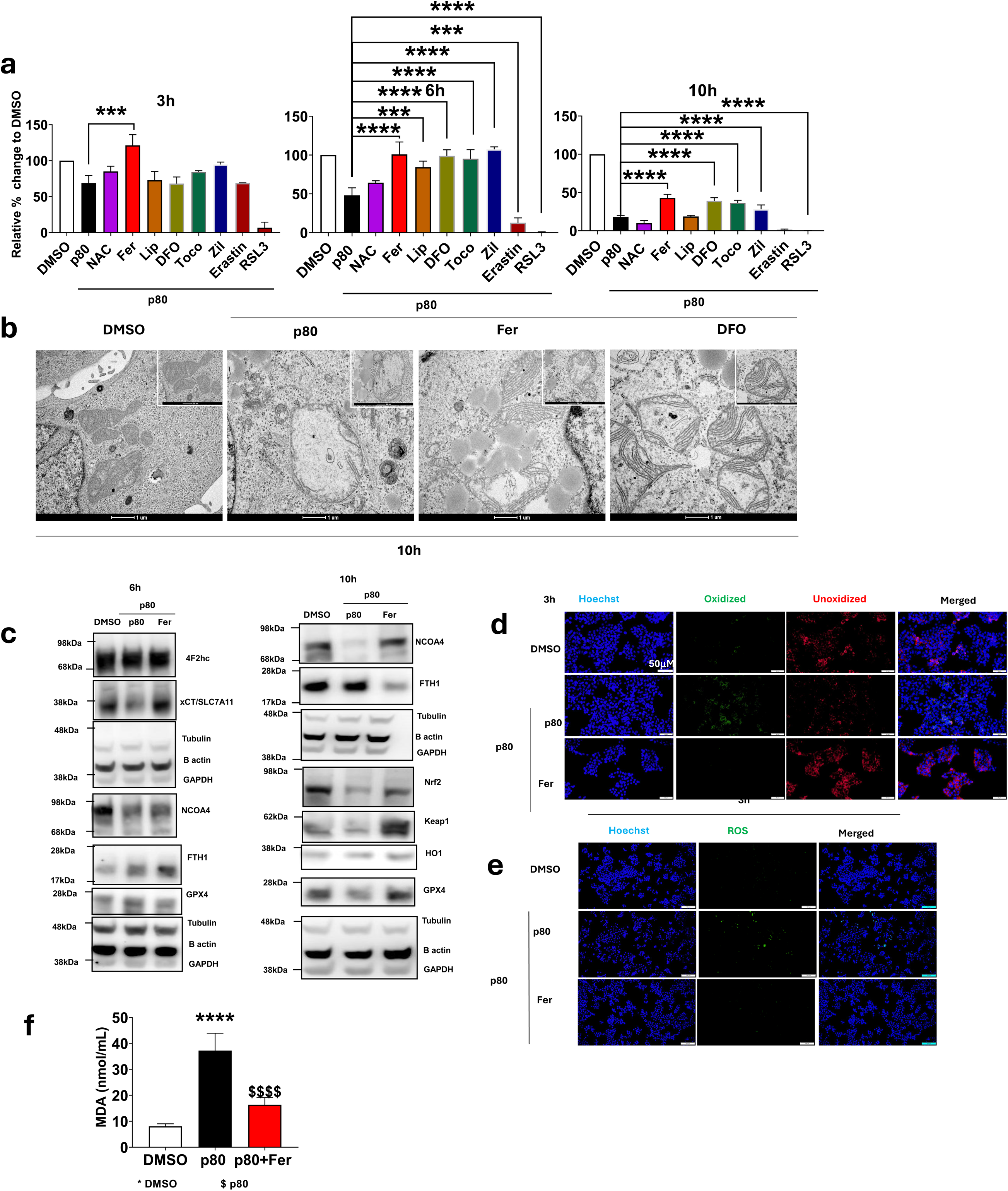
Ferroptosis inhibitors mitigate p80-induced mitochondrial damage and oxidative stress in HT29 cells and murine intestine. **(a)** Cell viability (CellTiter-Glo) at 3, 6, and 10 h after treatment with 0.25 % p80 and Ferroptosis inhibitors Liproxstatin-1 (Lip), Tocopherol (Toco), Deferoxamine (DFO), or Zileuton (Zil); DMSO as vehicle control. Y-axis = Viability (% of time-matched DMSO = 100%). **(b)** Transmission electron microscopy (TEM) of mitochondrial ultrastructure at 10 h after treatment with DMSO, p80, or p80 co-treated with Fer or DFO; scale bar, 1 µm. **(c)** Western blot analysis of ferroptosis markers (GPX4, SLC7A11, FTH1) and oxidative stress marker HO-1 at 10 h post p80 ± Fer; β-actin, GAPDH, and Tubulin used as loading/housekeeping controls. **(d, e)** Confocal microscopy showing lipid peroxidation (D, red fluorescence) and ROS production (E, green fluorescence) in HT29 cells at 3 h post-p80 and Fer-treatment; scale bars, 50 µm. **(f)** Malondialdehyde (MDA) in small-intestinal loops from live C57BL/6 female mice treated for 2 h with vehicle (DMSO in PBS), 1% polysorbate-80 (p80), or p80 + 10 µM Ferrostatin-1 (Fer). Mice per group: DMSO n=4; p80 n=5; p80+Fer n=5. Graphs show mean ± SD from three independent biological experiments. Statistical significance determined by one-way ANOVA with Tukey’s test (*P < 0.05; **P < 0.01; ***P < 0.001; ****P < 0.0001 versus control). (f) ****P < 0.0001 versus DMSO control and ^$$$$^ P < 0.0001 versus p80.

Based on these findings, Fer and DFO were selected for Transmission electron microscopy (TEM) studies (Figure 4b). TEM images at 10 hours revealed that p80 alone induced severe mitochondrial swelling and cristae disruption, which was partially alleviated upon co-treatment with Fer or DFO, suggesting that both iron chelation (DFO) and direct ferroptosis inhibition (Fer) can mitigate the mitochondrial damage triggered by p80.

Since similar effects were observed with various ferroptosis inhibitors in previous experiments, subsequent studies focused solely on Fer. At 10 hours, Western blot analysis showed that Fer co-treatment restored the p80-induced reduction of xCT/SLC7A11 and NCOA4 protein levels (Figure 4c). Moreover, Fer increased KEAP1 expression and led to modest elevations in GPX4 and Nrf2 compared to p80 alone, while HO-1 levels remained unchanged. This data further supports p80 targets ferroptosis to induce cell death in IECs.

Confocal microscopy images revealed a markedly reduction in p80-induced lipid peroxidation and ROS production after 3 hours co-treatment with Fer (Figures 4d,e), confirming p80-induced oxidative damage at is regulated by ferroptosis.

To further validate our *in vitro* findings to *in vivo* setting, the accumulation of malondialdehyde (MDA), a marker of lipid peroxidation, was assessed in small intestinal loops (Schwarzer et al., 2023) in living mice co-treated with p80 and Fer (Figure 4f). The results show a significant increase in MDA levels in the p80-treated group compared to the DMSO group, which was reduced upon co-treatment of p80 with Fer in the intestine (Figure 4f). This data agrees with the *in vitro* results and further supports p80 activation of ferroptosis in the intestine in living animals.

### Polysorbate 80 (p80) induces lipid droplet accumulation and lipid metabolism reprogramming in HT29 cells

RNA-seq analysis identified an upregulation in genes associated with lipid metabolism induced by p80 after 2 hours (Figure 1d). Further network analysis was conducted to elucidate the intricate interactions between genes implicated in lipid metabolism and ferroptosis pathways following p80 treatment. The Molecular Complex Detection (MCODE) network analysis of lipid metabolism genes, identified 26 upregulated genes associated to p80-induced ferroptosis, highlighting p80’s extensive regulatory impact on lipid metabolic processes (Figure 5a). The MCODE algorithm was employed to examine specific genes altered by p80 connecting lipid metabolism and ferroptosis (Figure 5b). ACSL1-ASCL5 were linked to p80-induced ferroptosis as well as ACSL1. Furthermore, LPCAT3’s role in membrane lipid composition alteration was linked to p80’s effect (Figure 5b). Further analysis of genes shared between ferroptosis, and lipid metabolism pathways identified upregulation of genes including *ACSL1, LPCAT3, ACSL4*, and *FDFT1*, which are crucial for fatty acid metabolism, activation, and lipid biosynthesis (Yang et al., 2022) (Table 3). Key regulators like *HMGCR* (involved in cholesterol biosynthesis regulation (Lee et al., 2024) were also significantly altered, reflecting an early shift in lipid homeostasis and oxidative stress (Table 3). *ACSL4* and *LPCAT3* enrich cellular membranes with polyunsaturated fatty acids (PUFAs) that are the primary substrates for lethal peroxidation (Doll et al., 2017). At 6 hours, further upregulation of genes associated with lipid metabolism was observed including *ACACA,* a pivotal regulator of lipid metabolism (Dong et al., 2024), *INSIG1*, linked to lipid biosynthesis (Wang et al., 2016) and *LPIN1*, involved in sterol regulation (Table 4) (Ishimoto et al., 2009). These results highlight possible profound alteration in lipid metabolism following p80 exposure.

**Figure 5:**
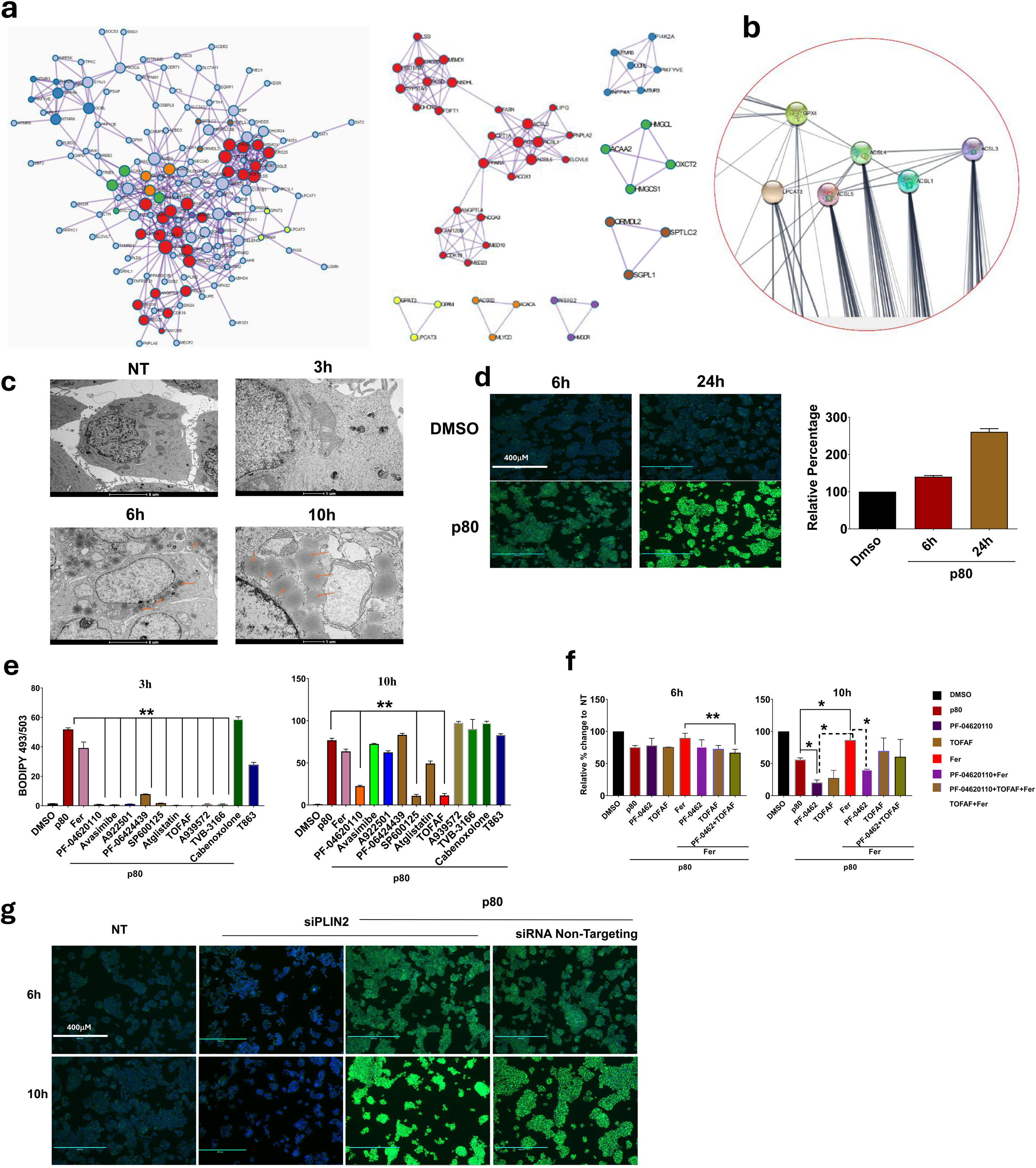
p80 induces lipid droplet formation and activates lipid metabolism pathways in HT29 cells, mitigated by inhibitors. **(a)** RNA-seq network of differentially expressed genes in HT29 cells treated with 0.25% p80, highlighting upregulated (red) and downregulated (blue) genes in lipid metabolism and ferroptosis pathways. **(b)** Gene Ontology (GO) enrichment of lipid metabolism–related pathways activated by 0.25% p80. **(c)** Transmission electron microscopy (TEM) of HT29 cells over 10 h of 0.25% p80 treatment; orange arrows mark lipid droplets. Scale bar: 1 µm. **(d)** Confocal images and quantification of BODIPY-stained lipid droplets in HT29 cells treated with 0.25% p80 for 6 h and 24 h; scale bar: 400 µm. Left, relative per centage BODIPY-stained area, data are mean ± SEM (n = 3); one-way ANOVA with Tukey’s post hoc (**p < 0.01; ***p < 0.001 versus NT). **(e)** Neutral lipid droplets by BODIPY 493/503 flow cytometry after 0.25% p80 for 3 h (left) and 10 h (right) ± inhibitors (Avasimibe, PF-04620110 [DGAT1], A-922500 [DGAT1], PF-06424439 [DGAT2], Atglistatin [ATGL], TOFA [ACC], A939572 [SCD1], TVB-3166 [FASN], Carbenoxolone, T863 [DGAT1]) or Ferrostatin-1 (Fer).Y-axis: % BODIPY-high cells (AF488-A, 530/30; live singlets). For more detail go to Supplementary figure 6. **(f)** CellTiter-Glo viability at 6 h and 10 h in HT29 cells treated with 0.25% p80 ± PF-04620110, (DGAT1), TOFA (ACC), or Ferrostatin-1 (Fer), alone or combined. X-axis: treatment groups (NT, p80, p80+PF, p80+TOFA, p80+Fer, combos). Y-axis: Viability (% of time-matched NT = 100%). Mean ± SEM (n = 3); one-way ANOVA with Tukey’s (*p < 0.05; **p < 0.01; ***p < 0.001). **(g)** Confocal BODIPY images of HT29 cells transfected with PLIN2 siRNA versus non-targeting control after 0.25% p80 treatment; scale bar: 400 µm. Representative of three experiments.

**Table 3.**
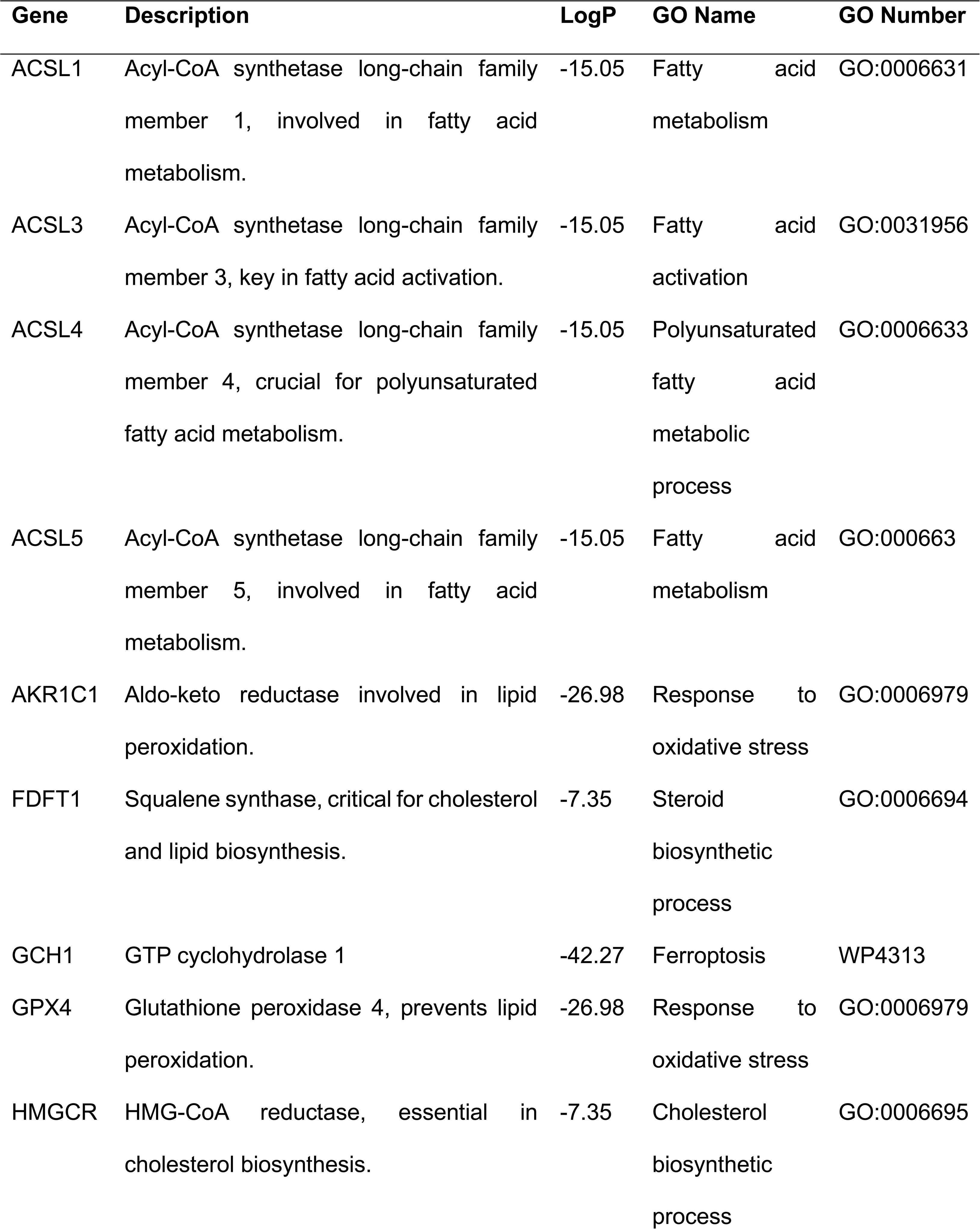
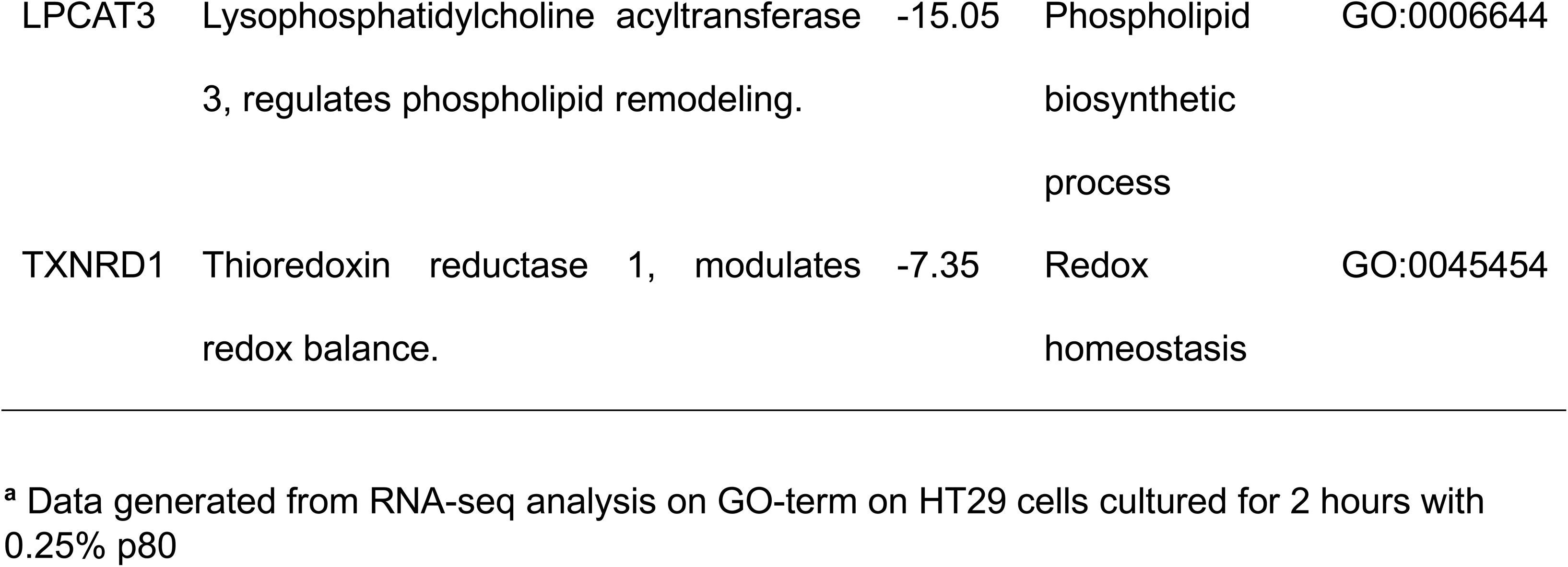
RNAseq analysis of most important upregulated genes of 2h shared ferroptosis and lipid metabolism pathways^a^.

**Table 4.** RNAseq analysis of most important upregulated lipid metabolism genes at 6.

| Gene | Description | LogP | GO Name | GO Number |
| --- | --- | --- | --- | --- |
| **ACACA** | Acetyl-CoA carboxylase alpha | -37.93 | Metabolism of lipids | R-HSA-556833 |
| **CYP1B1** | Cytochrome P450 family 1 subfamily B member 1 | -25.79 | Cellular response to lipid | GO:0071396 |
| **HMGCR** | 3-Hydroxy-3-Methylglutaryl-CoA Reductase | -13.51 | Cholesterol biosynthetic process | GO:0006695 |
| **HMGCS1** | 3-Hydroxy-3-Methylglutaryl-CoA Synthase 1 | -13.51 | Sterol biosynthetic process | GO:0016126 |
| **INSIG1** | Insulin-induced gene 1 | -33.49 | Lipid biosynthetic process | GO:0008610 |
| **LPIN1** | Lipin 1 | -10.89 | Sterol regulatory element-binding | WP1982 |
| **MVD** | Mevalonate Diphosphate Decarboxylase | -19.19 | Cholesterol metabolic process | GO:0008203 |
| **PLA2G6** | Phospholipase A2 Group VI | -13.11 | Lipid catabolic process | GO:0016042 |
| **PNPLA2** | Patatin-Like Phospholipase Domain Containing 2 | -10.03 | Glycerolipid biosynthetic process | GO:0045017 |
| **PPARD** | Peroxisome proliferator-activated receptor delta | -21.72 | Alcohol metabolic process | GO:0006066 |

<sup>a</sup> Data generated from RNA-seq analysis on GO-term on HT29 cells cultured for 6 hours with 0.25% p80

To validate these findings and examine the impact of p80 on lipid metabolism and its role in ferroptosis, lipid droplet formation and the activation of lipid metabolic pathways was examined in HT29 cells treated with 0.25% p80. Transmission electron microscopy revealed a progressive time-dependent accumulation of lipid droplets up to 10 hours p80-exposure compared to non-treated (NT) control cells (Figure 5c). Using BODIPY™ 493/503 staining, lipid droplets were found to be increased in both number and size over 24 hours of p80 treatment (Figure 5d).

To further investigate the pathways involved in lipid metabolism during p80-induced lipid droplet formation, small molecule inhibitors targeting key enzymes in lipid metabolic pathways were employed. Flow cytometry analysis using BODIPY staining demonstrated that all tested inhibitors, including Avasimibe, PF-04620110, A922501, SP600125, Atglistatin, A939572, TVB-3166, Carbexolone, and T863, effectively reduced lipid droplet formation at 3 hours post-p80 treatment (Figure 5e). However, by 10 hours, two inhibitors namely PF-04620110, a diacylglycerol acyltransferase-1 (DGAT-1) inhibitor, and 5-tetradecyl-oxy-2-furoic acid (TOFAF), an Acetyl-CoA-carboxylase-alpha (ACCA) inhibitor, presented a sustained and significant mitigating effect on p80-induced lipid droplet accumulation (Figure 5e).

Based on the findings on the MCODE network analysis (Figure 5a), a combination of lipid and ferroptosis inhibitors (DGAT-1, TOFAF and Fer, respectively) was examined for the potential recovery of p80-induced cell death phenotype. Inhibition of lipid metabolism either by the individual inhibitors or by their combinations with Fer did not recover the cell death phenotype at 6- or 10-hour time-points (Figure 5f), although it reduced lipid droplet formation.

Perilipin 2 (PLIN2) is a lipid droplet-associated protein critical for maintaining lipid droplet stability and regulating lipid metabolism under stress conditions (Xu et al., 2019). It has been implicated in cellular responses to oxidative stress and ferroptosis, where its upregulation is associated with lipid peroxidation and tumour progression (Bezawork-Geleta et al., 2022). To examine whether PLIN2 was involved in p80-induced ferroptosis, siRNA-mediated knockdown of PLIN2 was performed in HT29 cells. Confocal microscopy images at 6- and 10-hours post-treatment (Figure 5g) revealed no significant reduction in lipid droplet formation compared to non-targeting siRNA-treated cells, nor did it mitigate the p80-induced cell death at 24 hours (Supplementary Figure 4a). Similarly, the p80-induced cell death was not recovered in PLIN2 siRNA transfected cells over a 24-hour time frame (Supplementary figure 4a).

qPCR analysis of 12 lipid-related genes showed that *PLIN2* was significantly upregulated under p80 treatment, while *SEIPIN, PLIN3, SCD1, and LPCAT1/2* were downregulated (Supplementary Figure 4b). These findings suggest that PLIN2 alone is not essential for p80-induced lipid droplet accumulation or cell death, indicating the involvement of additional lipid regulatory pathways.

### Polysorbate 80 (p80) induces significant alterations in lipidomic profile in HT29 cells

To elucidate the intricate lipid pathways and networks modulated by p80 and their possible regulatory roles in the observed phenotype, a lipidomic analysis was undertaken. The analyses were focused on triglycerides, mono- and poly unsaturated fatty acids (MUFAs and PUFAs), lipids known to be associated with lipid droplet accumulation and ferroptosis (Bezawork-Geleta et al., 2022). A kinetic study was undertaken and Principal Component Analysis (PCA) scores plot reveals Principal Component 1 (PC1) capturing 75.3% of the variance, markedly distinguishing between the p80 treated and NT groups while PC2 explains 11.2% of the variation; with some overlap between the two groups (Supplementary Figure 5a). Next, a heatmap analysis of differentially expressed triglycerides at 10 hours post-treatment was performed (Figure 6a), showing significant clustering alterations in triglyceride composition between p80-treated and non-treated (NT) cells. Addition of the identified lipid metabolism inhibitors TOFAF, and DGAT revealed significantly reduced p80-induced TG levels. Co-treatment of cells with TOFAF, DGAT together with Fer did not appear to further reduce TG levels, thus indicating inhibitors targeting lipid enzymes effectively mitigate p80-induced TG alterations (Figure 5b). PCA analysis (Supplementary Figure 5b) corroborated these findings, showing distinct clustering of treatment groups and the influence of inhibitors in restoring lipid metabolic balance in the context of p80-induced stress.

**Figure 6:**
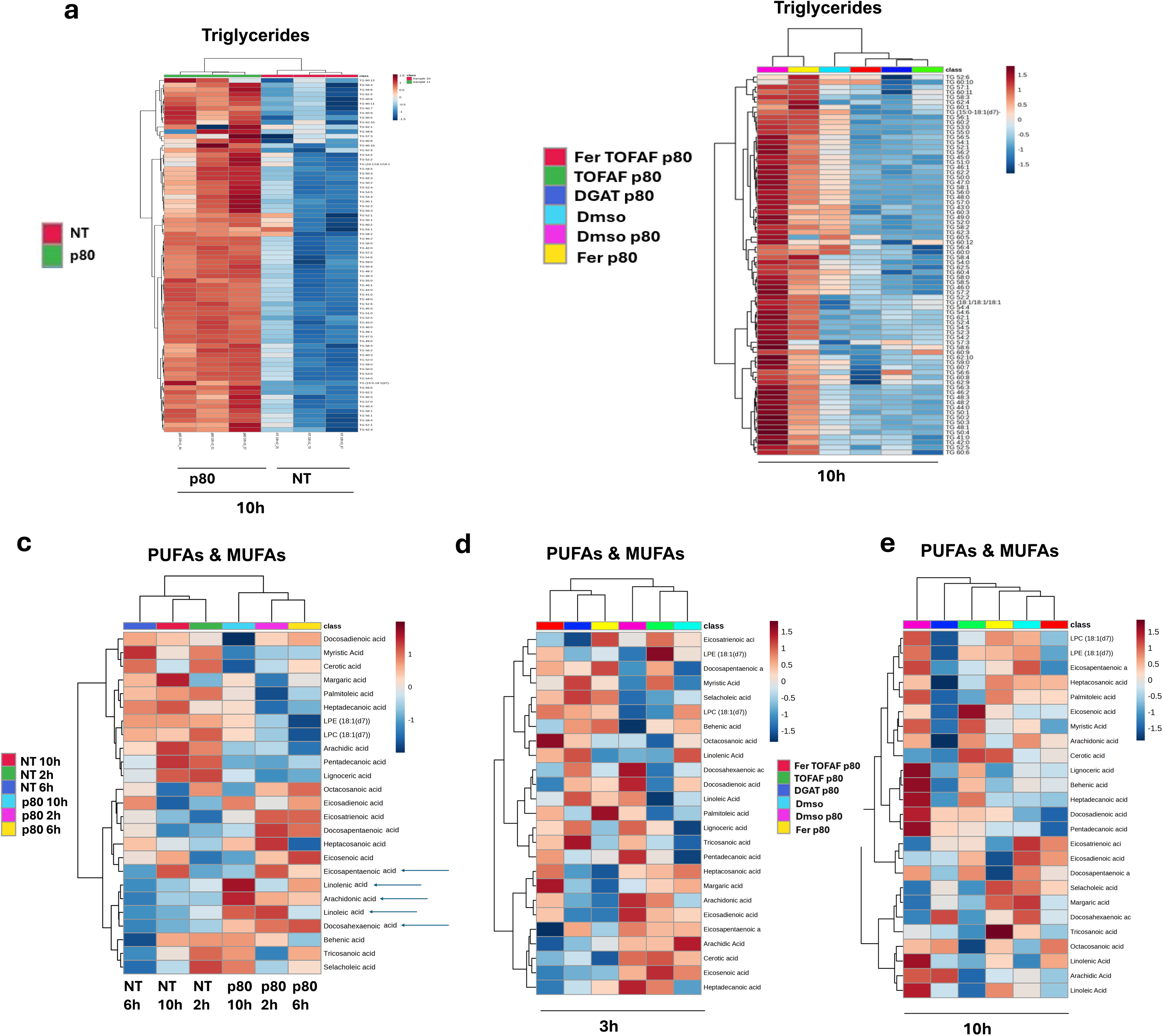
Lipidomic analysis reveals p80-induced changes in gene expression and fatty acid composition in HT29 cells. **(a)** Heatmap of differentially expressed triglycerides in HT29 cells at 10 hours post-treatment with 0.25% p80 compared to non-treated (NT) controls. **(b)** Heatmap showing the effects of Ferrostatin-1 (Fer), TOFA, and a DGAT inhibitor on 0.25% p80-induced triglyceride accumulation at 10 hours post-treatment. **(c)** Heatmap of polyunsaturated fatty acids (PUFAs) and monounsaturated fatty acids (MUFAs) at 2, 6, and 10 hours post-treatment with 0.25% p80. **(d, e)** Heatmaps of PUFAs and MUFAs highlighting the impact of lipid metabolism inhibitors at 3 hours **(d)** and 10 hours (e) on 0.25% p80-induced lipid alterations. NT = non-treated control. Values are log2 fold-changes vs time-matched NT from aligned, median-normalized LC–MS intensities. A 16:0 sphingomyelin internal standard monitored extraction and instrument stability. Red = increase; blue = decrease.

A kinetic and PCA analysis on PUFAs and MUFAs shows that PC1 accounts for 21.1% and PC2 for 17.4% of the variance in these lipids showing a distinct clustering between p80-treated and NT samples over time (Supplementary Figure 5c). Heatmap analysis further revealed substantial changes in the profile of these lipids, with markedly elevated levels of key PUFAs, including linoleic acid, arachidonic acid, and eicosapentaenoic acid, in the p80-treated cells, particularly at the 10 hours timepoint (Figure 5c). Treatment with TOFAF and DGAT either alone or in combination with Fer revealed that p80-induced arachidonic, eicosadienoic, eicosapentaenoic docosadienoic, linoleic and linolenic acids are reduced upon Fer, DGAT, and TOFAF individual treatments (Figure 6d,e). There is no enhanced reduction in the combination treatments at 3- and 10-hour timepoints. In contrast, docosapentaenoic acid is paradoxically higher across all inhibitor treatments at 3 hour timepoint. p80-induced SFAs and MUFAs show a general decrease under inhibitor treatments, with reductions in heptacosanoic, heptadecanoic, and pentadecanoic acids, while increasing myristic acid compared to p80 treatment (Figure 6d,e). Myristic acid, which previously increased under p80, now is reduced in the DGAT, Fer, and their combination-treated cells. The earlier increase in palmitoleic acid induced by inhibitors is not sustained, except by Fer. Pentadecanoic acid continues to rise under p80 but is decreased by Fer and DGAT treatments. Finally, selacholeic/nervonic acid, which is reduced by p80, increases with Fer and TOFAF+Fer treatments. Tricosanoic acid shows a notable increase only with Fer treatment, highlighting the specific and varied impacts of these treatments on fatty acid profiles at 10 hours.

Docosahexaenoic was decreased at 3h and 10h (Figure 6 d,e). This might indicate that while p80 initially disrupts docosahexaenoic acid levels, the inhibitors are effective in preserving its presence, potentially offering protective effects against ferroptosis induced by p80 (Mortensen et al., 2023). These findings indicate a time-dependent changes in PUFAs and MUFAs, with accumulation of ferroptosis-associated PUFAs and reduction in MUFAs. A summary on the effect on p80-induced PUFAs, MUFAs and SFA after 10 hours inhibitor treatment is presented in Table 5.

**Table 5.**
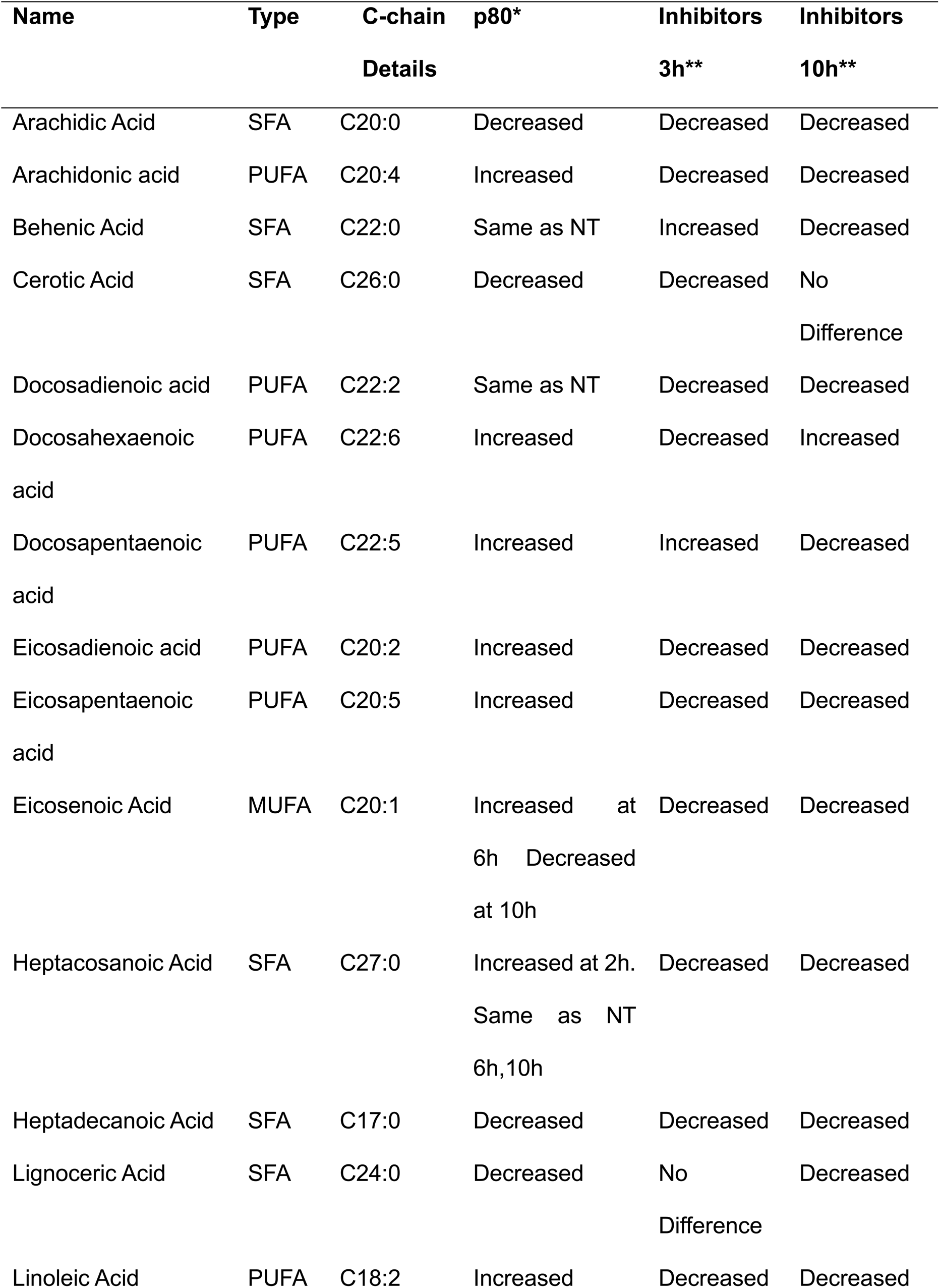
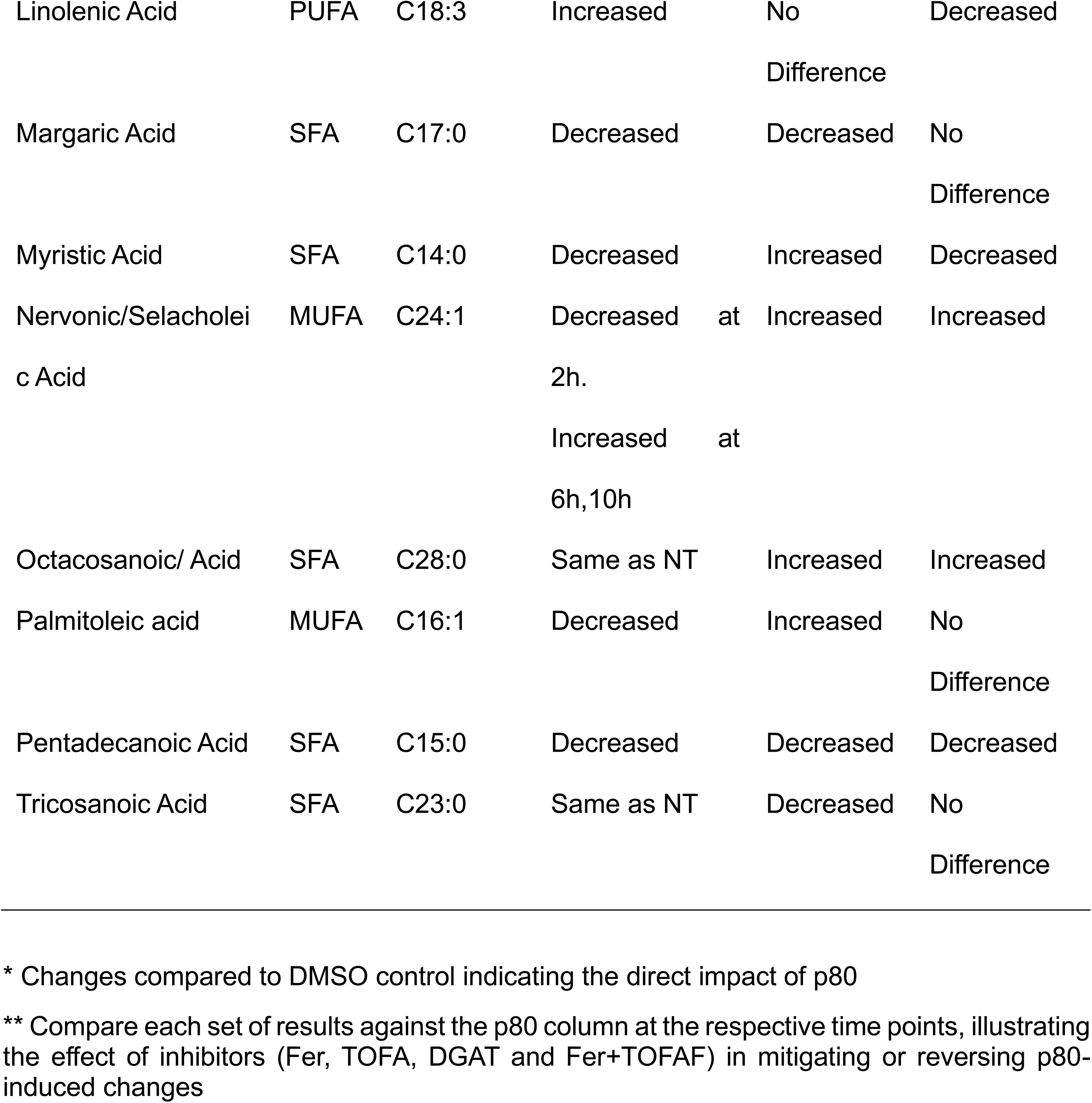
Summary on lipid and ferroptosis inhibitors’ effect on p80-induced PUFAs and MUFAs in epithelial cells.

## DISCUSSION

The increasing use of food additives such as emulsifiers like polysorbate-80 (p80) in processed foods has raised significant concerns about their potential negative impact on human health (Monteiro et al., 2019; Chassaing et al., 2017). Preclinical studies showed a negative impact of p80 and carboxymethylcellulose on the microbiota leading to worsening in several chronic conditions including inflammatory bowel disease (Chassaing et al., 2015). A mechanistic understanding on the influence of these emulsifiers on the host, particularly intestinal epithelial cells is lacking. This study shows that p80 negatively affects intestinal epithelial cells responses by altering lipid metabolism, oxidative stress, iron and lipid droplet accumulation, mitochondrial function and structure, and cell death via ferroptosis. This data provide a new mechanistic understanding on the direct influence of certain emulsifiers, p80, on intestinal epithelial cell biology, which can potentially contribute to increase incidence in chronic diseases.

p80 is widely used for its emulsifying properties, which stabilize and improve the consistency of a wide range of products (Food Safety Commission of Japan, 2007), but its potential to disrupt gut health and cellular homeostasis has brought it under scrutiny (Chassaing et al., 2015; Schwartzberg et al., 2018). Findings from this study support a role for p80 as a contributor to ferroptosis and link its effects to the pathogenesis of chronic diseases such as inflammatory bowel disease and metabolic syndrome (Hou et al., 2011; Friedmann-Angeli et al., 2014). Ferroptosis is primarily regulated by GPX4, a critical enzyme that mitigates lipid peroxidation and cellular damage by reducing hydroperoxides to alcohols (Friedmann-Angeli et al., 2014). Previous studies have shown that markers associated with ferroptosis including GPX4, iron accumulation, were markedly altered in colon biopsies and IECs of patients with ulcerative colitis (UC) and in animal models of IBD (Xu et al., 2020). Treatment with ferroptosis inhibitors including Ferrostatin-1 reversed colitis and ferroptosis markers (such as FTH1 and GPX4) in animals with colitis (Chen et al.,2020). In addition, IECs from patients with UC underwent ferrotoptic cell death via NFkB ferroptotic inhibition (Xu et al., 2020). Alterations to mitochondrial structure, ROS and mitochondrial iron accumulation and direct oxidative damage to lipids was uncovered by p80. However, alterations on GPX4 protein caused by p80 treatment over time were subtle, despite its significant reduction in the RNASeq data set, indicating p80-induces cell death in gut epithelial cells is driven by GPX4 independent mechanism(s) (Gao et al., 2019; Su et al., 2019). Other genes reported as associated with the GPX4-independent pathway (*FSP1, DHODH and GCH1*) (Escuder-Rodríguez et al., 2024.) were not altered by p80, further supporting the notion that p80-induced ferroptosis in gut cells is independent of GPX4 (Gao et al., 2019; Su et al., 2019). p80 rapidly impacts mitochondria inducing a significant reduction in mitochondrial membrane potential. This is reminiscent of cystine-starvation induced ferroptosis, where mitochondria become hyperpolarized and fuel lipid peroxidation (Gao et al., 2019). The structural changes observed in mitochondria, such as disrupted cristae and mitochondrial shrinkage, are hallmarks of ferroptosis and indicated by the p80 disruption of mitochondrial integrity (Gao et al., 2019). A hyperpolarized mitochondrion produces more superoxide as the proton gradient builds without adequate ATP synthesis. By 3 h, overt mitochondrial damage was observed by TEM p80-treated cells exhibited swollen mitochondria with disrupted cristae, a morphology often seen in ferroptotic cells (Gao et al., 2019). This mitochondrial dysfunction contributes to increased ROS production, which can result in exacerbated oxidative stress and lipid peroxidation, contributing to a feedback loop that amplifies ferroptotic cell death (Stockwell et al., 2017; Gao et al., 2019).

The addition of emulsifiers like p80 into drug formulations adds another level of complexity to their potential impact on the health of cells and the microbiota (Maier et al., 2018; Chassaing et al., 2015). p80’s emulsifying properties enhance the solubility and bioavailability of hydrophobic drugs, making it a widely used component in drug delivery systems (Ravichandran et al., 2021; U.S. Food and Drug Administration, 2023). Several studies have demonstrated that p80 can cause microbial alterations, increase intestinal permeability and exacerbate inflammation, highlighting the need for careful evaluation of its use in therapeutic formulations (Chassaing et al., 2015; Li et al., 2020). Maier et al. (2018) identified several marketed drugs that affect the gut microbiota, and when focusing on drugs used in the treatment of inflammatory bowel disease, their analysis revealed that five such drugs have microbiota-impacting properties. Among these, at least four: Infliximab (U.S. Food and Drug Administration, 2013), Ustekinumab (Drugs.com, 2023), Entyvio (European Medicines Agency, 2023) and Stelara (Janssen, 2018), contain p80 in their formulation. In addition, it is plausible that all members of the polysorbate family (p20, p60 and p80) have the potential to be toxic owing to their similar emulsifying properties (Food Safety Commission of Japan, 2007; Schwartzberg et al., 2018). A recent human trial revealed no impact of 6 days oral consumption of p80 on intestinal membrane fluidity or intestinal membrane disruption in healthy volunteers (Metry <u>et</u> al., 2022). Some oncolytic drugs formulations containing p80 have shown an effect on the drugs’ distribution and elimination upon i.v. administration, which can result in an increased systemic exposure and reduced release of the drug (Schwartzberg et al.,2018). Other side effects reported by p80-containing formulations include hypersensitivity, nonallergic anaphylaxis, rash and injection- and infusion site adverse events (ISAEs) as well as some clinical cases of renal and liver toxicity (Schwartzberg et al.,2018). Several IBD-associated biologicals e.g. anti-TNFs, anti-integrins, anti-IL12/23, contain p80 in their formulations, but it is currently unknown whether ISAEs from these biologics is caused due to p80. The findings on p80’s effects on gut epithelial cells, particularly in individuals with underlying vulnerabilities such as IBD, are yet to be explored, raising potential concerns about the balance between functional benefits over safety in drug formulations.

Our findings integrate well with the recent work by Ogulur et al. (2023), who showed that p80 can disrupt the gut barrier and alter gene expression in human intestinal cells and organoids (Ogulur et al., 2023). They observed upregulation of ferroptosis-related genes (like SLC7A11 and PRNP) with emulsifier exposure, which they interpreted as a protective response against toxicity (Ogulur et al., 2023).

In addition to its effects on mitochondrial function, p80 exposure contributed to the accumulation of lipid droplets in the cells, suggesting an adaptive cellular response to sequester excess lipids and prevent lipotoxicity. The upregulation of genes involved in lipid droplet formation, such as PLIN2, further supports this adaptive response (Zadoorian et al., 2023). Interestingly addition of small molecules regulating autophagy, or in this context lipophagy, which is relevant for the degradation of excess lipid droplets, or the knock-down of PLIN2, did not revert the p80-induced cell death phenotype.

The increase in TAGs likely represents the cell sequestering excess fatty acids into inert storage forms (droplets) as a protective strategy against lipotoxicity. By esterifying free PUFAs into TAGs, cells can temporarily reduce the substrate available for peroxidation in membranes. Indeed, higher levels of certain species containing C18:2 (linoleic) and C20:4 (arachidonic) fatty acids were detected in p80-treated samples, which may indicate PUFA rerouting into storage. However, despite this response, it appears to be insufficient to fully prevent lipid peroxidation and ferroptotic cell death. The lipid droplets themselves can become peroxidized over time, and the continued presence of p80 likely feeds more fatty acids into the system than can be safely stored or detoxified, overwhelming the cell’s protective capacity and ultimately triggering ferroptosis.

Although the lipid metabolism inhibitors, TOFAF (an acetyl-CoA carboxylase inhibitor) and DGAT, effectively reduced p80-induced lipid droplet formation, they did not fully recover the cytotoxic effects regulated by p80, not even in the presence of ferroptosis inhibitors, indicating the involvement of multiple interconnected pathways, including those related to mitochondrial dysfunction, oxidative stress, and disrupted autophagy (Su et al., 2019).

This study revealed that p80-induced lipid droplets significantly accumulated triglycerides and PUFAs such as arachidonic acid (AA) and docosahexaenoic acid (DHA) (Shah et al., 2018). PUFAs are integral to numerous cellular processes, but their susceptibility to lipid peroxidation under oxidative stress creates a pro-ferroptotic environment (Dixon et al., 2012). This implies that phospholipases (for example, cytosolic phospholipase A₂s) are active, cleaving fatty acyl chains from membrane phospholipids, a response to get rid of damaged lipids (Dixon et al., 2012). However, free PUFAs in the cytosol can be double-edged: while removing them from membranes can be protective, those free fatty acids can themselves undergo auto-oxidation or be oxygenated by enzymes e.g., free AA can be oxygenated by 5-LOX or 15-LOX into peroxides. The net effect detected in the lipidomic analysis is that free PUFA levels are increased but did not prevent cell death, implying that any detoxification via phospholipase activity was outpaced by the continued generation of new peroxides (Chen et al., 2021)

A recent report shows that a westernised diet, high in PUFAs, especially AA, induces a pro-inflammatory cytokine response in gut epithelial cells which is regulated by GPX4 and results in ileal inflammation reminiscent of CD in GPX4-transgenic mice (Mayr et al., 2020). In contrast, p80 barely changed GPX4 levels/expression in IECs (this study) and p80 induced IL-8 gene expression within 2 hours of p80-treatment of HT29 cells (Saiz et al., 2021). Further examination on HMGB1, a damage-associated molecular pattern molecules, released by ferroptotic cells (Lee et al., 2024) was not detected in p80-treated gut epithelial cells (not shown). This data indicates that p80-induced ferroptosis in gut epithelial cells, under the conditions examined in this study, do not trigger an inflammatory response.

The cholesterol and isoprenoid lipid pathways were also perturbed by p80, which may indirectly sensitize cells to ferroptosis. The transcriptomics data showed strong upregulation of cholesterol biosynthetic enzymes (*HMGCR*, *HMGCS1, MVK,* etc. (Madison., 2016) at 6 h, suggesting cellular cholesterol was depleted or dysfunctional. Based on the collected data, it is postulated that p80, being a surfactant, may extract cholesterol from plasma membranes or interfere with its distribution, prompting the cell to ramp up synthesis. It is noteworthy that the *INSIG1* gene, which encodes a cholesterol feedback regulator, was highly up-regulated at 6 h, and *LDLR* (LDL receptor) was also induced, reinforcing that the cells sensed low cholesterol. Thus, p80 creates a scenario akin to cholesterol starvation of membranes, inadvertently favoring lipid peroxidation processes.

In the gastrointestinal tract, emulsifiers can disrupt the mucosal barrier, leading to increased intestinal permeability (“leaky gut”) and enabling the translocation of microbial products that trigger inflammation (Chassaing et al., 2015). Systemically, these effects may extend to metabolic tissues, where emulsifiers could interfere with lipid signalling pathways, exacerbate insulin resistance, and contribute to adipose tissue inflammation with broader implications for systemic health, including metabolic and cardiovascular diseases (Rizzello et al., 2019; Monteiro et al., 2019). Additionally, the emulsifiers’ potential to alter gut microbiota composition could influence host immunity and metabolic homeostasis, further amplifying their physiological impact (Vindigni et al., 2016). Chronic exposure to p80 could potentiate these risks by exacerbating oxidative stress and promoting dysregulated lipid metabolism, particularly in tissues with high lipid turnover such as the liver and adipose tissue (Narula et al., 2021; Shah et al., 2018).

The broader implications of these findings are significant, particularly for individuals susceptible for or suffering of several chronic conditions including IBD, metabolic syndrome, and early-onset cancers. Oxidative stress and lipid dysregulation are key drivers of these conditions, and the ability of dietary emulsifiers like p80 to amplify these processes warrants stricter regulatory scrutiny (Chassaing et al., 2015; Narula et al., 2021). Findings in pilot studies in healthy volunteers consuming another emulsifier, CMC, added to an emulsifier free diet for 10 days, revealed some abdominal discomfort and microbiota alterations in the volunteers (Chassaing et al., 2021) Other studies where patients with quiescent UC were fed carrageenan for 7 days revealed no major impact on systemic inflammation or gastrointestinal symptoms (Laatikainen et al., 2023). In a randomised, double-blind, placebo-controlled, re-supplementation trial in patients with mild/moderately active CD fed a low emulsifier content diet for 8 weeks resulted in reduced clinical activity index with no impact on fecal calprotectin (Bancil, et al., 2025), while in a double-blinded, randomized, controlled feeding study, with patients with ileal active CD on a low emulsifier diet for 4 weeks showed no effect on intestinal inflammation. Overall, these findings show the need to further evaluate the broader implications of emulsifiers on human health, particularly in vulnerable populations with pre-existing metabolic or inflammatory conditions.

## LIMITATIONS

Limitations with this study include the use of 2D *in vitro* cell systems and the lack of other cell types such as immune cells and the microbiota, which may not fully reflect the *in vivo* conditions. Although studies on effects of p80 in intestinal loops was conducted, the study lacks long-term exposure data. Further research is needed to explore microbiome interactions, genetic and disease variability, and *in vivo* validation of these findings.

## CONCLUSION

In summary, this study has identified mechanisms by which a well-utilised emulsifier, p80, disrupts lipid metabolism, induces ferroptosis, and compromise’s mitochondrial function in intestinal epithelial cells, advancing our understanding of the negative influences on a dietary emulsifier on cellular health. By addressing ferroptosis and lipid metabolism pathways, this work highlights the importance of mitigating the risks associated with p80 exposure through targeted research, regulatory measures, and therapeutic innovations.

## Acknowledgements

The authors like to acknowledge Ms Tiina O’Neill and Dr Dimitri Scholz, UCD, for the support and discussions on processing, analysis and imaging of cells on transmission electron microscopy; Dr Aine Fanning for training and support to GS-G at the start of the project; Dr Ciaran Lee for training and support to GS-G on siRNA experiments; Dr Eileen Ryan for advise to GS-G on lipidomic analysis. Flow cytometry analysis was performed at the APC Microbiome Ireland Flow Cytometry Platform located at University College Cork.

## Funding

This research was conducted with the financial support of Science Foundation Ireland (SFI) under grant number SFI/12/RC/2273-P2. GS-G is a recipient of a Government of Ireland Postgraduate Scholarship (grant GOIPG/2019/4528). MS, DS and TH acknowledge support from Talking microbes – understanding microbial interactions within One Health framework (CZ.02.01.01/00/22_008/0004597) from the Ministry of Education, Youth and Sports of the Czech Republic.

## Contributors

GS-G, SAJ and SM: conception of the work; GS-G and SM: design of the work; GS-G, RS and NH: experiments with organoids and western blotting; GS-G and GC: confocal microscopy; GS-G, SMa and JOS: Seahorse studies; GS-G, KQ, SAJ: lipidomics set up, optimisation and analysis; CM and GS-G: analysis of RNASeq data; GS-G and TC: flow cytometry; GS-G: siRNA experiments; DS, TH, MS: intestinal loop experiments; GS-G: drafted the manuscript; GS-G, and SM: critical revision of the work for important intellectual content. All authors made significant contributions to the paper and approved the final version. SM is the lead contact and guarantor.

## STAR Methods

### Key Resources

#### Table Key resources table 1

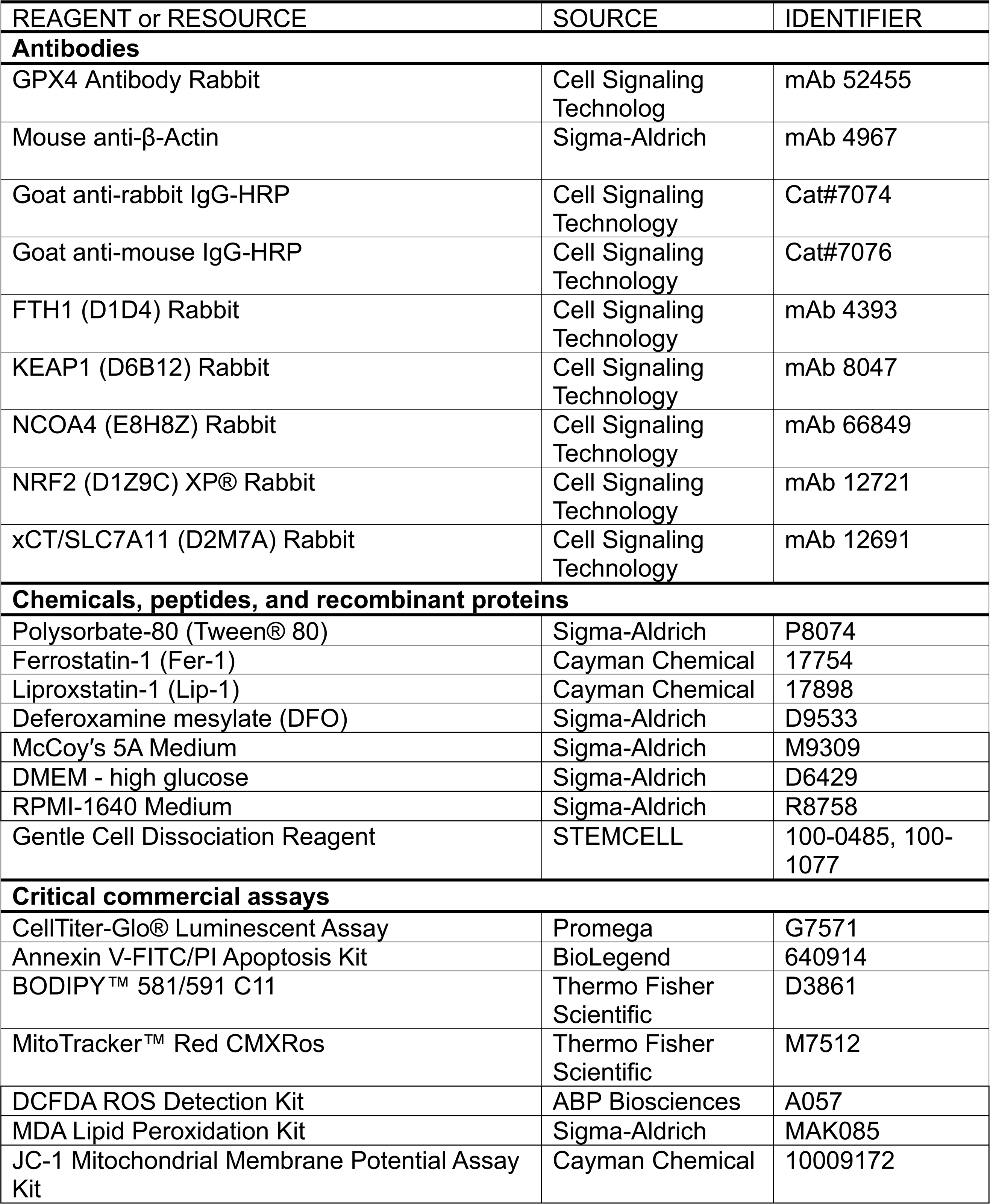
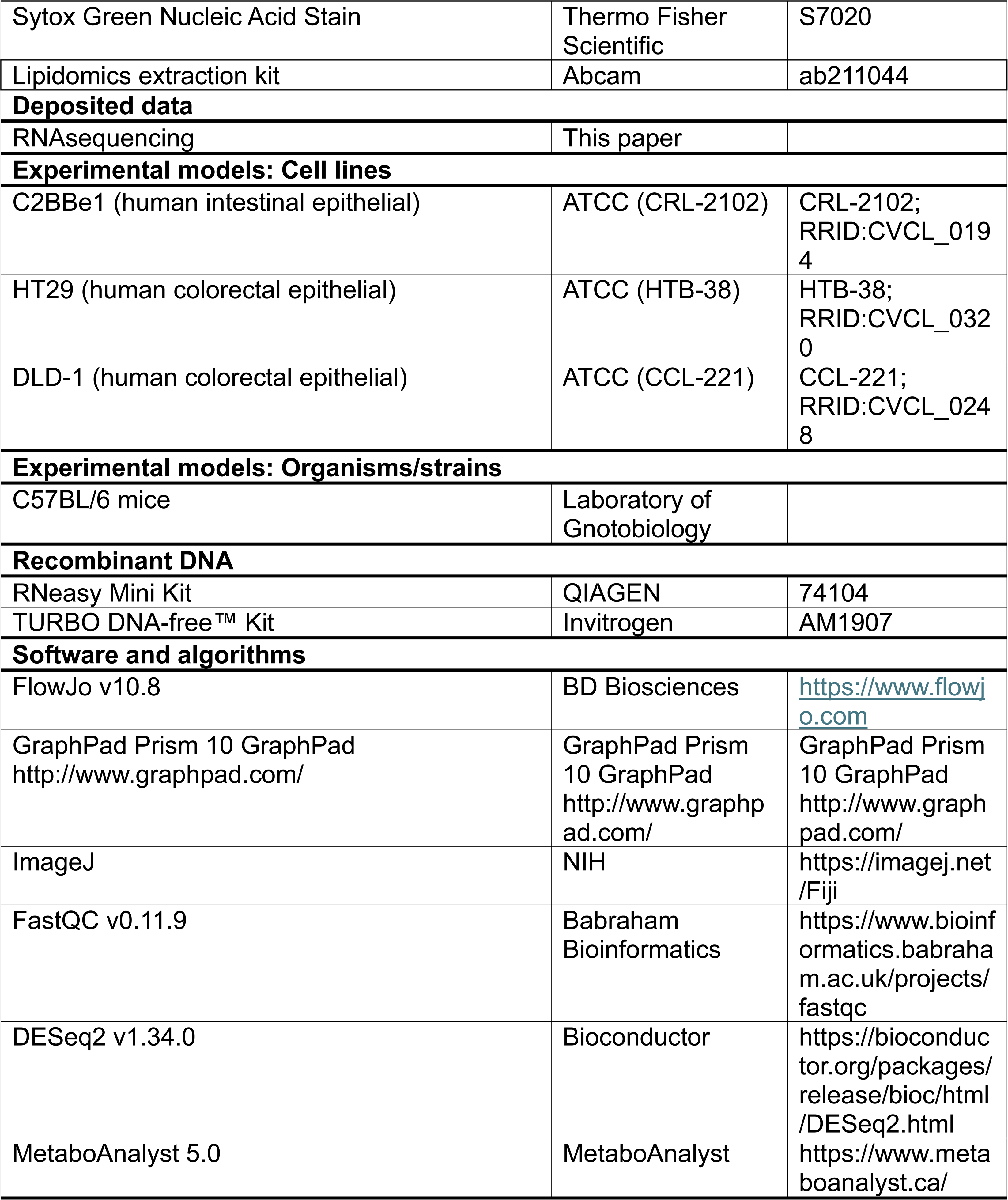

| REAGENT or RESOURCE | SOURCE | IDENTIFIER |
| --- | --- | --- |
| **Antibodies** |  |  |
| GPX4 Antibody Rabbit | Cell Signaling Technology | mAb 52455 |
| Mouse anti-β-Actin | Sigma-Aldrich | mAb 4967 |
| Goat anti-rabbit IgG-HRP | Cell Signaling Technology | Cat#7074 |
| Goat anti-mouse IgG-HRP | Cell Signaling Technology | Cat#7076 |
| FTH1 (D1D4) Rabbit | Cell Signaling Technology | mAb 4393 |
| KEAP1 (D6B12) Rabbit | Cell Signaling Technology | mAb 8047 |
| NCOA4 (E8H8Z) Rabbit | Cell Signaling Technology | mAb 66849 |
| NRF2 (D1Z9C) XP® Rabbit | Cell Signaling Technology | mAb 12721 |
| xCT/SLC7A11 (D2M7A) Rabbit | Cell Signaling Technology | mAb 12691 |
| **Chemicals, peptides, and recombinant proteins** |  |  |
| Polysorbate-80 (Tween® 80) | Sigma-Aldrich | P8074 |
| Ferostatin-1 (Fer-1) | Cayman Chemical | 17754 |
| Liproxstatin-1 (Lip-1) | Cayman Chemical | 17898 |
| Deferoxamine mesylate (DFO) | Sigma-Aldrich | D9533 |
| McCoy's 5A Medium | Sigma-Aldrich | M9309 |
| DMEM - high glucose | Sigma-Aldrich | D6429 |
| RPMI-1640 Medium | Sigma-Aldrich | R8758 |
| Gentle Cell Dissociation Reagent | STEMCELL | 100-0485, 100-1077 |
| **Critical commercial assays** |  |  |
| CellTiter-Glo® Luminescent Assay | Promega | G7571 |
| Annexin V-FITC/PI Apoptosis Kit | BioLegend | 640914 |
| BODIPY™ 581/591 C11 | Thermo Fisher Scientific | D3861 |
| MitoTracker™ Red CMXRos | Thermo Fisher Scientific | M7512 |
| DCFDA ROS Detection Kit | ABP Biosciences | A057 |
| MDA Lipid Peroxidation Kit | Sigma-Aldrich | MAK085 |
| JC-1 Mitochondrial Membrane Potential Assay Kit | Cayman Chemical | 10009172 |
| Sytox Green Nucleic Acid Stain | Thermo Fisher Scientific | S7020 |
| Lipidomics extraction kit | Abcam | ab211044 |
| **Deposited data** |  |  |
| RNAsequencing | This paper |  |
| **Experimental models: Cell lines** |  |  |
| C2BBE1 (human intestinal epithelial) | ATCC (CRL-2102) | CRL-2102;<br>RRID:CVCL_0194 |
| HT29 (human colorectal epithelial) | ATCC (HTB-38) | HTB-38;<br>RRID:CVCL_0320 |
| DLD-1 (human colorectal epithelial) | ATCC (CCL-221) | CCL-221;<br>RRID:CVCL_0248 |
| **Experimental models: Organisms/strains** |  |  |
| C57BL/6 mice | Laboratory of Gnotobiology |  |
| **Recombinant DNA** |  |  |
| RNeasy Mini Kit | QIAGEN | 74104 |
| TURBO DNA-free™ Kit | Invitrogen | AM1907 |
| **Software and algorithms** |  |  |
| FlowJo v10.8 | BD Biosciences | https://www.flowjo.com |
| GraphPad Prism 10 GraphPad<br>http://www.graphpad.com/ | GraphPad Prism 10 GraphPad<br>http://www.graphpad.com/ | GraphPad Prism 10 GraphPad<br>http://www.graphpad.com/ |
| ImageJ | NIH | https://imagej.net/Fiji |
| FastQC v0.11.9 | Babraham Bioinformatics | https://www.bioinformatics.babraham.ac.uk/projects/fastqc |
| DESeq2 v1.34.0 | Bioconductor | https://bioconductor.org/packages/release/bioc/html/DESeq2.html |
| MetaboAnalyst 5.0 | MetaboAnalyst | https://www.metaboanalyst.ca/ |

#### Lead Contact

Further information and requests for resources and reagents should be directed to and will be fulfilled by the Lead Contact, Dr. Silvia Melgar.

#### Materials Availability

All unique materials generated in this study are available from the Lead Contact.

#### Data and Code Availability

The RNA sequencing and lipidomics data will be deposit on NCBI GEO and MetaboLights. All other data supporting the findings of this study are available from the corresponding author upon reasonable request. No restrictions apply to the availability of code.

### EXPERIMENTAL MODEL AND STUDY PARTICIPANT DETAILS

#### Jejunal loops

Specific pathogen-free C57Bl/6 mice were kept in Laboratory of Gnotobiology in IVC cages (Tecniplast, Italy), exposed to 12:12 hours light-dark cycles, supplied with filtered non-chlorinated tap water, and fed ad libitum mouse breeding diet V1124-300 (Ssniff Spezialdiäten GmbH) sterilized by irradiation (∼25 kGy, Bioster, Czech Republic). Female C57Bl/6 mice were anesthetized by the intraperitoneal administration of a ketamine/xylazine mixture. Mouse abdomen was shaved and laparotomy was performed. Two jejunal loops 2 cm long per mouse were created with nylon ligatures. 10 uM Ferrostatin-1 (Merck Life Science, Czech Republic) and 1% Polysorbate 80 dissolved in DMSO/PBS or 1% Polysorbate 80 in DMSO/PBS were injected into jejunal loops through a gauge needle (100 μl). In control loops only 100 ul of DMSO/PBS was injected. After surgery, the jejunal loops was returned to the abdominal cavity, which was then sutured, and the mouse was placed on heated pad (37 °C). After 2 hours, the mouse was euthanized, the abdominal cavity was re-opened, and the jejunal loop was removed. Each loop was divided into 4 parts which were 1) snap-frozen in liquid N2, 2) embeded in Tissue-Tek® O.C.T. ™ (Life M, Czech Republic) and frozen in liquid N2 or 3) placed in the RNAlater (Merck Life Science, Czech Republic) at 4°C overnight and subsequently frozen at −80°C. The experiments were approved by the Animal Care Committee of the Czech Academy of Sciences (protocol n. 18/2019) and were in accordance with the EU Guide for the Care and Use of Laboratory Animals.

#### Cell Lines

Human intestinal epithelial cell lines HT29 (ATCC HTB-38), C2BBe1 (ATCC CRL-2102) and DLD-1 (ATCC CCL-221), were obtained from the American Type Culture Collection (ATCC). HT29 and C2BBe1 cells were maintained in McCoy’s 5A medium (Gibco, Thermo Fisher Scientific), while DLD-1, all supplemented with 10% fetal bovine serum (FBS; Sigma-Aldrich), 100 IU/ml penicillin, and 0.1 mg/mL streptomycin (Gibco, Thermo Fisher Scientific). Cells were authenticated by short tandem repeat profiling by the supplier (ATCC) and periodically tested for *Mycoplasma* using PCR (abm Good to Go™ kit, cat #G238). Cells were incubated at 37 °C in a humidified atmosphere of 5 % CO_2_ and used for experiments between passages 7 and 15. Cells were passaged at approximately 80% confluency using 0.25% Trypsin-EDTA (Gibco, Thermo Fisher Scientific).

#### Small Intestinal Organoids

##### Crypt Isolation and Culture

Crypts were isolated from the distal half of the small intestine of C57BL/6 mice (bred in the Biological Sciences Unit, University College Cork, Cork) following established protocols (Sato et al., 2013) with some changes. Briefly, the small intestine was carefully excised and immediately placed into ice-cold phosphate-buffered saline (PBS). The intestine was gently flushed to remove luminal contents, opened longitudinally, and thoroughly washed in fresh PBS. The cleaned intestinal tissue was then sectioned to facilitate subsequent crypt isolation. Crypts were embedded in BME-2 (Amsbio) and cultured in Intesticult™ Organoid Growth Medium (STEMCELL Technologies). Organoids were maintained at 37 °C in 5% CO_2_, with media changes every 2 days. Organoid viability was measured using the CellTiter-Glo 3D Cell Viability Assay (Promega) and Sytox Green stain (Thermo Fisher Scientific).

### METHOD DETAILS

#### Emulsifier Preparation

Polysorbate 80 (p80; Sigma-Aldrich, P1754), carboxymethylcellulose (CMC; Sigma-Aldrich, C5678), were prepared as 1% stock solutions in the respective culture media.

#### Assessment of Cell Viability, Apoptosis, and Proliferation

Cell viability, apoptosis, and proliferation were assessed using multiple assays. Cell viability was evaluated with the CellTiter-Blue® Cell Viability Assay (Promega) and the CellTiter-Glo® Luminescent Cell Viability Assay (Promega), both conducted according to the manufacturer’s instructions, with luminescence measured using a GloMax® Discover Microplate Reader (Promega). Apoptosis was measured using the Caspase-Glo® 3/7 Assay (Promega), following the manufacturer’s protocol, and luminescence was similarly read on a GloMax® Discover Microplate Reader.

#### Western Blot

Protein expression levels following various treatments were analyzed by western blotting as previously reported (Saiz-Gonzalo et al., 2021). Briefly, cells were treated as indicated, washed with cold PBS, and lysed using RIPA buffer supplemented with protease and phosphatase inhibitors. Protein concentrations were determined using a BCA assay. Equal protein amounts (40 µg) were denatured and separated on 4–12% Bis-Tris gels, then transferred onto PVDF membranes. Membranes were blocked with 5% skim milk in TBST for 1 hour, incubated overnight at 4°C with primary antibodies, followed by HRP-conjugated secondary antibodies for 1 hour at room temperature. Bands were visualized with chemiluminescent HRP substrate and detected using a LAS-3000 Imager. Image analysis was performed using ImageJ without image modification.

#### RNA Isolation and Quantitative Reverse Transcription PCR (RT-qPCR)

Total RNA was isolated using the RNeasy Mini Kit (Qiagen) according to the manufacturer’s instructions. Cells were lysed directly in RLT buffer with 10 μL/mL β-mercaptoethanol (β-ME) (Sigma), and the lysates were homogenized, mixed with 70% ethanol, and passed through RNeasy spin columns. RNA was eluted in RNAse-free water and quantified using a NanoDrop spectrophotometer (Thermo Fisher Scientific). RNA integrity was assessed based on A260/A280 and A260/A230 ratios. Genomic DNA contamination was removed using the TURBO DNA-free™ Kit (Invitrogen). cDNA was synthesized from 500 ng of RNA using Transcriptor Reverse Transcriptase (Roche) with random primers, according to the manufacturer’s protocol.

Quantitative PCR was performed using SensiFAST™ Probe No-ROX Kit (Meridian Bioscience) on a LightCycler 480 (Roche). Gene expression was normalized to β-actin using the 2-ΔΔCt method (Livak and Schmittgen, 2001). All reactions were performed in at least duplicate, with appropriate positive and negative controls. Primers for RT-qPCR were designed using the Design Center Platform (Roche) and sequences are available upon request.

#### RNA Sequencing and Analysis

For transcriptomic analysis, total RNA was extracted from HT29 cells using the Qiagen RNeasy Mini Kit (74104). RNA quality and integrity were confirmed by Agilent Bioanalyzer 2100 (RIN ≥ 8.0). Raw sequence reads were assessed for quality with FastQC (v0.11.9) and trimmed for adapters using Trimmomatic (v0.39). Reads were aligned to the human reference genome (GRCh38) using the STAR aligner (v2.7.10a) with default parameters. FeatureCounts (Subread v2.0) was used to generate gene-level read counts based on GENCODE v35 annotations. Differential gene expression analysis was performed using the DESeq2 package (v1.34.0) in R (v4.2.1). Genes with an adjusted p-value < 0.05 and |log2 fold change| > 1 were considered significantly different. Gene set enrichment analysis (GSEA) was performed on normalized counts using the clusterProfiler package (Bioconductor) to identify enriched pathways.

#### Flow Cytometry

Cells were seeded in 6-well plates at a density of 1 × 10^6 cells per well and cultured overnight. Following treatment under the specified experimental conditions, cells were harvested by trypsinization, washed twice with cold PBS, and resuspended in staining buffer. Flow cytometric analysis was performed using a BD FACSCelesta™ flow cytometer (BD Biosciences) equipped with lasers suitable for FITC and PE detection. Data were collected for 10,000-20,000 events per sample and were analyzed using FlowJo™ software (BD Biosciences). The median fluorescence intensity (MFI) for each marker was determined and normalized to the control samples. Fluorescence compensation was applied to correct for spectral overlap between fluorophores.

#### MitoFerroGreen and BioTracker™ FerroOrange Live Cell Dye

##### Mitochondrial Iron Detection

To detect mitochondrial iron levels, HT29 cells were seeded in 6-well plates at a concentration of 1 × 10^6 cells per well. After reaching 80% confluence, cells were treated with the respective compounds for the indicated times. Following treatment, cells were washed twice with PBS and incubated with 1 μM MitoFerroGreen (Dojindo Laboratories) in complete medium for 30 minutes at 37 °C in a humidified incubator. After staining, cells were washed twice with PBS to remove excess dye and harvested by trypsinization. The cells were then resuspended in PBS and immediately analyzed by flow cytometry using a BD FACSCelesta™. MitoFerroGreen fluorescence was detected using an excitation wavelength of 488 nm and emission at 530 nm. The data were processed using FlowJo™ software, and the mean fluorescence intensity (MFI) was calculated. The MFI values were normalized to untreated controls to quantify mitochondrial iron levels.

##### Cytosolic Iron Detection

Cytosolic iron levels were assessed using BioTracker™ FerroOrange Live Cell Dye (Sigma-Aldrich). Cells were seeded in 6-well plates at a density of 1 × 10^6 cells per well and treated as per experimental requirements. After treatment, cells were washed twice with PBS and incubated with 1 μM FerroOrange in complete medium for 30 minutes at 37 °C. Post-incubation, cells were washed with PBS, harvested by trypsinization, and resuspended in PBS. Fluorescence imaging was performed using an Olympus FV3000 confocal microscope, with excitation at 561 nm and emission collected at 580 nm. Images were captured and analyzed using the associated software, and the fluorescence intensity was quantified to assess cytosolic iron levels. Results were normalized to control cells to provide a relative measure of cytosolic iron accumulation.

#### MitoTracker Staining and JC-1 Mitochondrial Membrane Potential Assay

##### MitoTracker Staining

To evaluate mitochondrial integrity, cells were seeded in 6-well plates at a density of 1 × 10^6 cells per well and allowed to adhere overnight. After treatment, cells were washed twice with PBS and incubated with 100 nM MitoTracker™ Red CMXRos (Invitrogen) in complete medium for 30 minutes at 37°C in the dark. Following staining, cells were washed again with PBS to remove unbound dye and fixed with 4% paraformaldehyde for 15 minutes at room temperature. Fixed cells were washed, resuspended in PBS, and visualized using an Olympus FV3000 confocal microscope. Fluorescence was detected with excitation at 579 nm and emission at 599 nm. Images were analyzed using ImageJ software to quantify the fluorescence intensity, which correlates with the mitochondrial mass and membrane potential.

##### JC-1 Mitochondrial Membrane Potential Assay

Mitochondrial membrane potential was measured using the JC-1 Mitochondrial Membrane Potential Assay Kit (Cayman Chemical). Cells were seeded in 6-well plates at a density of 1 × 10^6 cells per well and treated as required. Post-treatment, cells were washed twice with PBS and incubated with JC-1 dye at a final concentration of 2 μM in complete medium for 20 minutes at 37°C. After staining, cells were washed twice with PBS and analyzed by flow cytometry using a BD FACSCelesta™. The fluorescence of JC-1 monomers was measured with excitation at 488 nm and emission at 530 nm, while the fluorescence of JC-1 aggregates was measured with excitation at 561 nm and emission at 590 nm. The ratio of red (aggregates) to green (monomers) fluorescence was calculated to assess mitochondrial membrane potential. The data were analyzed using FlowJo™ software, and results were normalized to control samples.

#### Lipid Peroxidation and Reactive Oxygen Species (ROS) Detection

##### Lipid Peroxidation

Lipid peroxidation levels were measured using the BODIPY™ 581/591 C11 probe (Invitrogen), which selectively detects lipid peroxides in cellular membranes. Cells were seeded in 6-well plates at a density of 1 × 10^6 cells per well and treated with experimental compounds as specified. Post-treatment, cells were washed twice with PBS and incubated with 2 μM BODIPY C11 in complete medium for 30 minutes at 37°C. After staining, cells were washed with PBS, harvested by trypsinization, and resuspended in PBS. Flow cytometry was performed using a BD FACSCelesta™ flow cytometer. The fluorescence of non-oxidized BODIPY C11 was detected with excitation at 488 nm and emission at 530 nm, while oxidized BODIPY C11 was detected with excitation at 565 nm and emission at 590 nm. The ratio of oxidized to non-oxidized fluorescence was calculated to assess the degree of lipid peroxidation. The results were normalized to untreated controls to quantify relative lipid ROS production.

##### ROS Detection

Reactive oxygen species (ROS) levels were assessed using the ROS Detection Kit (Sigma-Aldrich), which includes the H2DCFDA fluorescent probe. Cells were seeded in 6-well plates at a density of 1 × 10^6 cells per well and treated as described. Following treatment, cells were washed twice with PBS and incubated with 10 μM H2DCFDA in complete medium for 20 minutes at 37°C in the dark. After incubation, cells were washed with PBS, harvested by trypsinization, and resuspended in PBS. Flow cytometry analysis was conducted using a BD FACSCelesta™ flow cytometer with excitation at 488 nm and emission at 530 nm. The mean fluorescence intensity (MFI) was used to quantify intracellular ROS levels, and data were normalized to control cells.

##### Seahorse XF Mito Stress Test (OCR/ECAR)

Mitochondrial respiration and glycolysis were quantified on an Agilent Seahorse XFe24 Analyzer using the XF Cell Mito Stress Test. HT29 cells were seeded into XFe24 cell-culture microplates to reach ∼80–90% confluence on the assay day (5×10⁴ cells/well). Where indicated, cells were pre-exposed to 0.25% (v/v) polysorbate-80 (p80) or vehicle for the stated times. One hour before the run, growth medium was replaced with XF Base Medium supplemented with 10 mM glucose, 1 mM sodium pyruvate, and 2 mM L-glutamine (pH 7.4), and plates were incubated 60 min in a non-CO₂ incubator. Sensor cartridges were hydrated overnight and loaded with the following final in-well concentrations: oligomycin 1 µM, FCCP 0.5–1 µM (titrated; 1 µM used for HT29), and rotenone/antimycin A 0.5 µM each. Data were acquired with standard mix/wait/measure cycles (3/2/3 min): three baseline measurements, then sequential injections from ports A–C (oligomycin → FCCP → rotenone/antimycin A). OCR (pmol O₂·min⁻¹) and ECAR (mpH·min⁻¹) were computed in Wave; assay parameters (ATP-linked OCR, proton leak, maximal respiration, spare capacity, non-mitochondrial OCR) followed Agilent equations.

#### Microscopy

##### Nile Red and BODIPY™ Staining

Lipid droplets in HT29 cells were visualized using Nile Red (Sigma-Aldrich) and BODIPY™ 493/503 (Invitrogen) staining. Cells were seeded in 6-well plates at a density of 1 × 10^6 cells per well and treated under specified conditions. After treatment, cells were washed twice with PBS and stained with Nile Red (1 μg/mL) or BODIPY™ 493/503 (2 μM) in complete medium for 30 minutes at 37°C. Following staining, cells were washed with PBS and fixed with 4% paraformaldehyde for 15 minutes at room temperature. Fixed cells were washed again with PBS and mounted on slides using Fluoroshield™ mounting medium (Sigma-Aldrich). Fluorescence images were acquired using an Olympus BX43 fluorescence microscope with appropriate filter sets. Nile Red fluorescence was detected with excitation at 552 nm and emission at 636 nm, while BODIPY™ 493/503 fluorescence was detected with excitation at 493 nm and emission at 503 nm. The images were processed and analyzed using ImageJ software to quantify fluorescence intensity, representing the extent of lipid accumulation within the cells.

##### Sytox Green Staining

Cell death was assessed using Sytox Green Nucleic Acid Stain (Thermo Fisher Scientific). Cells were seeded in 96-well plates and treated with the desired experimental conditions. Following treatment, cells were incubated with 1 μM Sytox Green for 15 minutes at room temperature, protected from light. After incubation, the cells were washed with PBS to remove excess dye, and fluorescence was measured using a SpectraMax iD3 Microplate Reader (Molecular Devices) with an excitation/emission of 504/523 nm. The fluorescence intensity correlates with the number of dead cells, as Sytox Green selectively penetrates and stains the nucleic acids of cells with compromised membranes.

##### MDA Assay

Lipid peroxidation levels in mouse intestinal tissue samples were quantified using the MAK085 Lipid Peroxidation (MDA) Assay Kit (Sigma-Aldrich) as per manufacturer’s instructions. The assay is based on the reaction of malondialdehyde (MDA) with thiobarbituric acid (TBA) to form a colorimetric product, measured at 532 nm. MDA standards were prepared to generate a standard curve, and the concentrations of MDA in the tissue samples were determined accordingly. The results were expressed in nmol/mL and compared across DMSO (control), p80-treated, and p80 with Ferrostatin-1 (Fer) groups to evaluate oxidative stress.

#### Lipidomics & analysis

Lipids from p80-treated HT29 cells were extracted using a commercial lipid extraction kit (ab211044, Abcam, UK) and stored at −80 °C for lipidomic profiling. For LC–MS, 100 µL of the dried extract (starting material per sample) was reconstituted and diluted 1:3 (v/v) with 100% isopropanol (IPA) containing 16:0 sphingomyelin (SM; Avanti Polar Lipids/Merck) at 1:40 (v/v) of the internal-standard working solution. The internal standard (16:0 SM, Avanti Polar Lipids) was obtained from Merck. Samples were analyzed using a Waters Xevo™ G2-XS QTOF Mass Spectrometer with an ACQUITY™ UPLC system. 5 µL were injected in technical triplicate onto an ACQUITY UPLC CSH™ C18 column (100 × 2.1 mm, 1.7 μm; 55 °C, 400 μL/min). Mobile phases: A (acetonitrile/water, 60:40) and B (isopropanol/acetonitrile, 90:10), both with 10 mM ammonium formate and 0.1% formic acid. MS operated in ESI+ and ESI− modes (m/z 100–2000, 1 s/scan, 120 °C source, 550 °C desolvation, 900 L/h nitrogen, 2.0 kV/1.5 kV capillary, 30 V cone). Leucine enkephalin (m/z 556.2771/554.2615) was used for lock mass correction.

Data were processed with Skyline-daily (MacCoss Lab Software) using an in-house lipid library (Dr. Susan Joyce). Untargeted LC-MS data were aligned in Progenesis QI and analyzed in MetaboAnalyst 5.0 (median normalization, log transformation, multivariate analysis).

#### Statistical analysis

GraphPad Prism software was used to perform statistical analyses for all data sets except RNA Sequencing and lipidomics (see respective section for their analysis). Unless specified, the data is expressed as mean ± SEM. A p-value <0.05 was considered significant.

## Notes

### Competing Interest Statement

The authors have declared no competing interest.

