## Supplementary Figures for "The dietary emulsifier polysorbate-80 induces lipid accumulation and cell death in intestinal epithelial cells via ferroptosis"

### Supplementary Figure 1

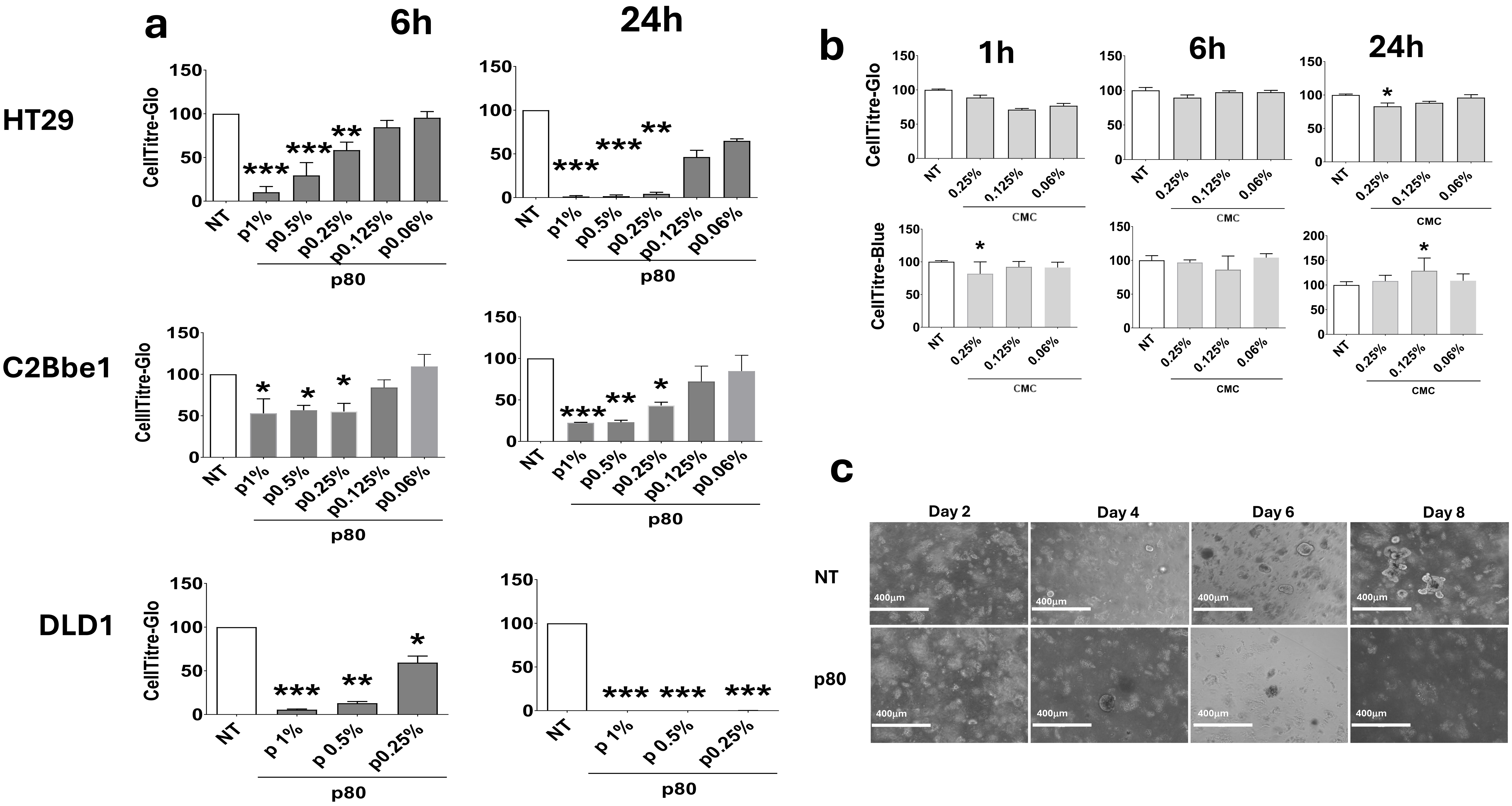

### Supplementary Figure 2

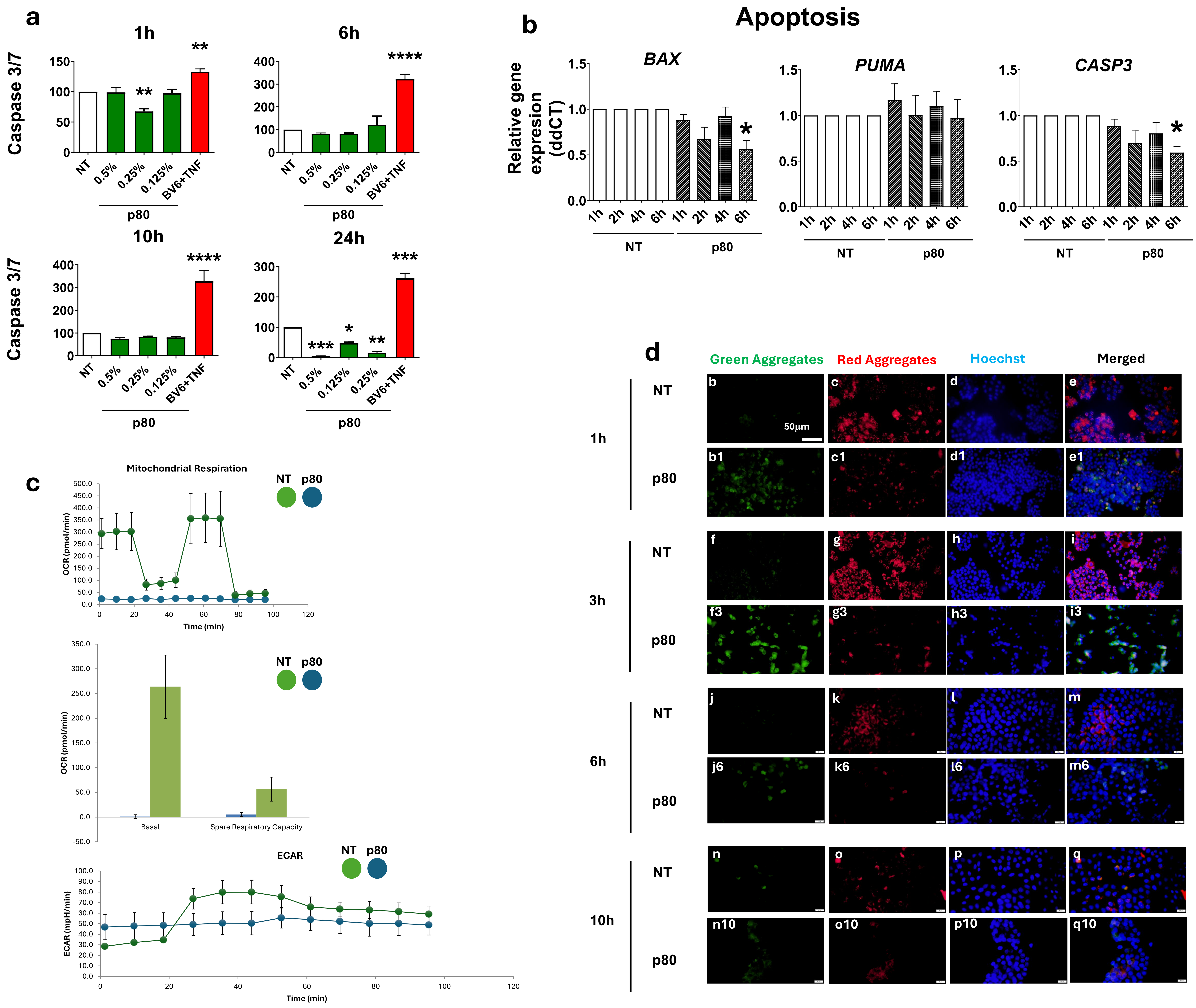

### Supplementary Figure 3

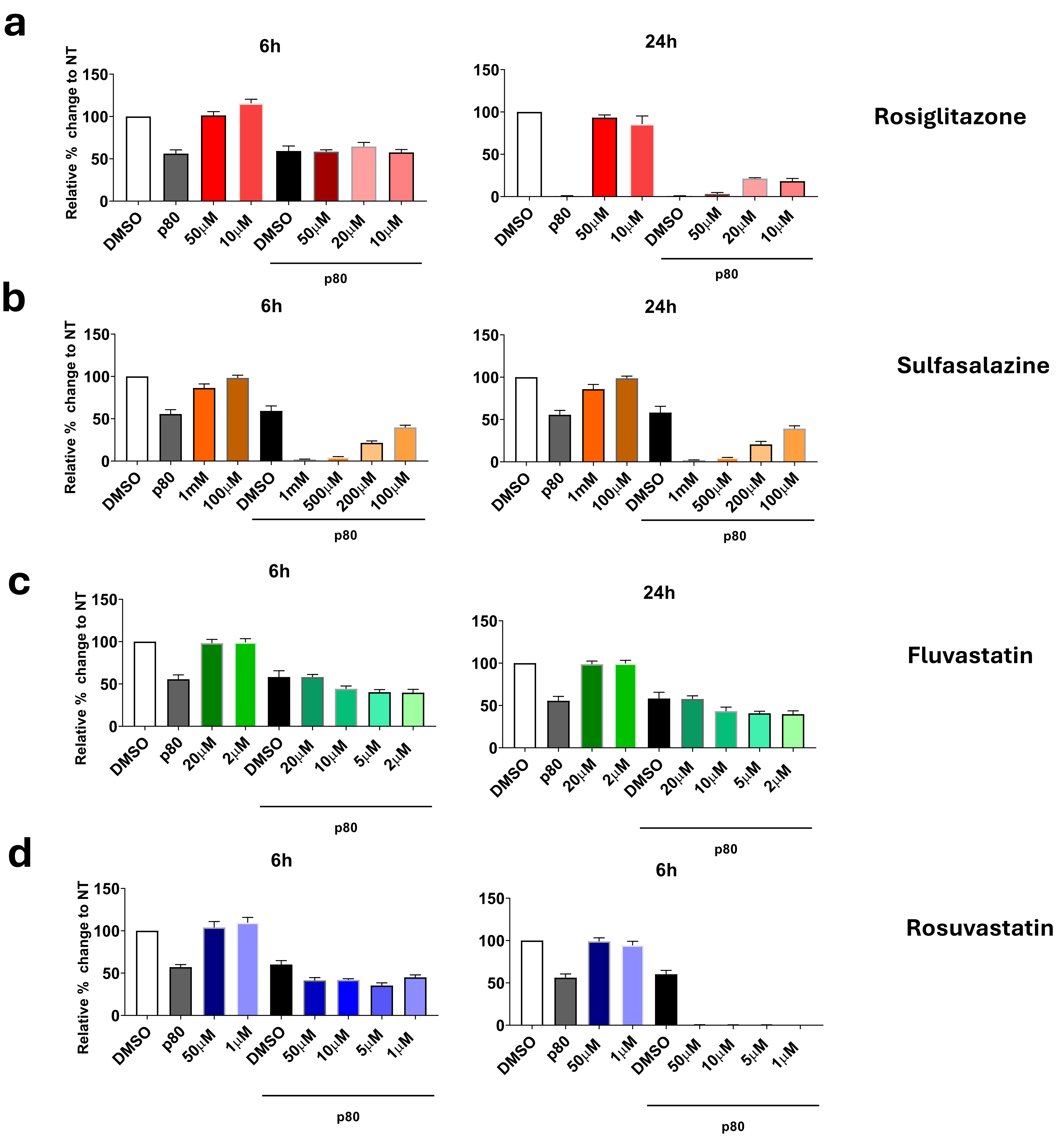

### Supplementary Figure 4

a

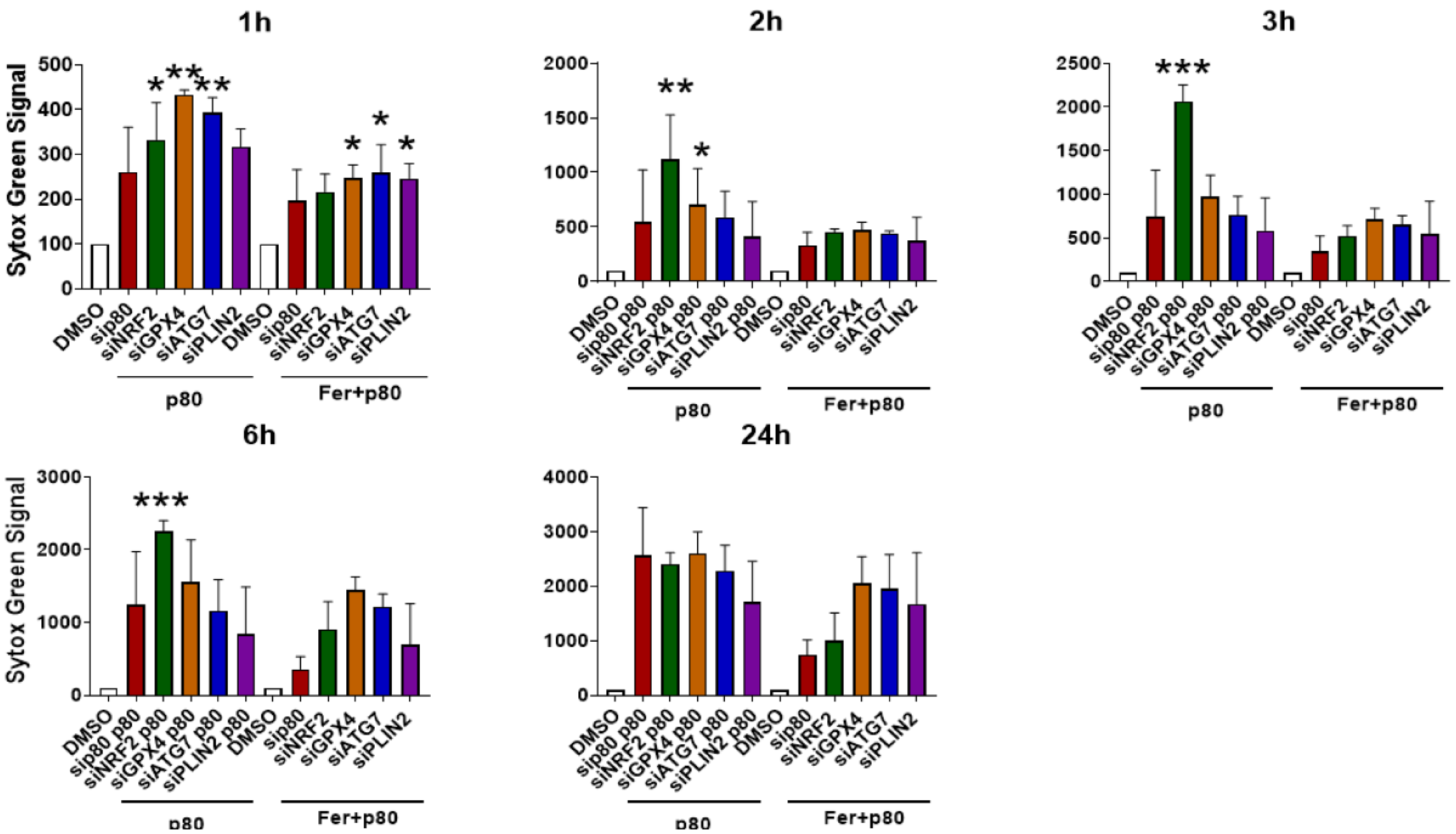


b

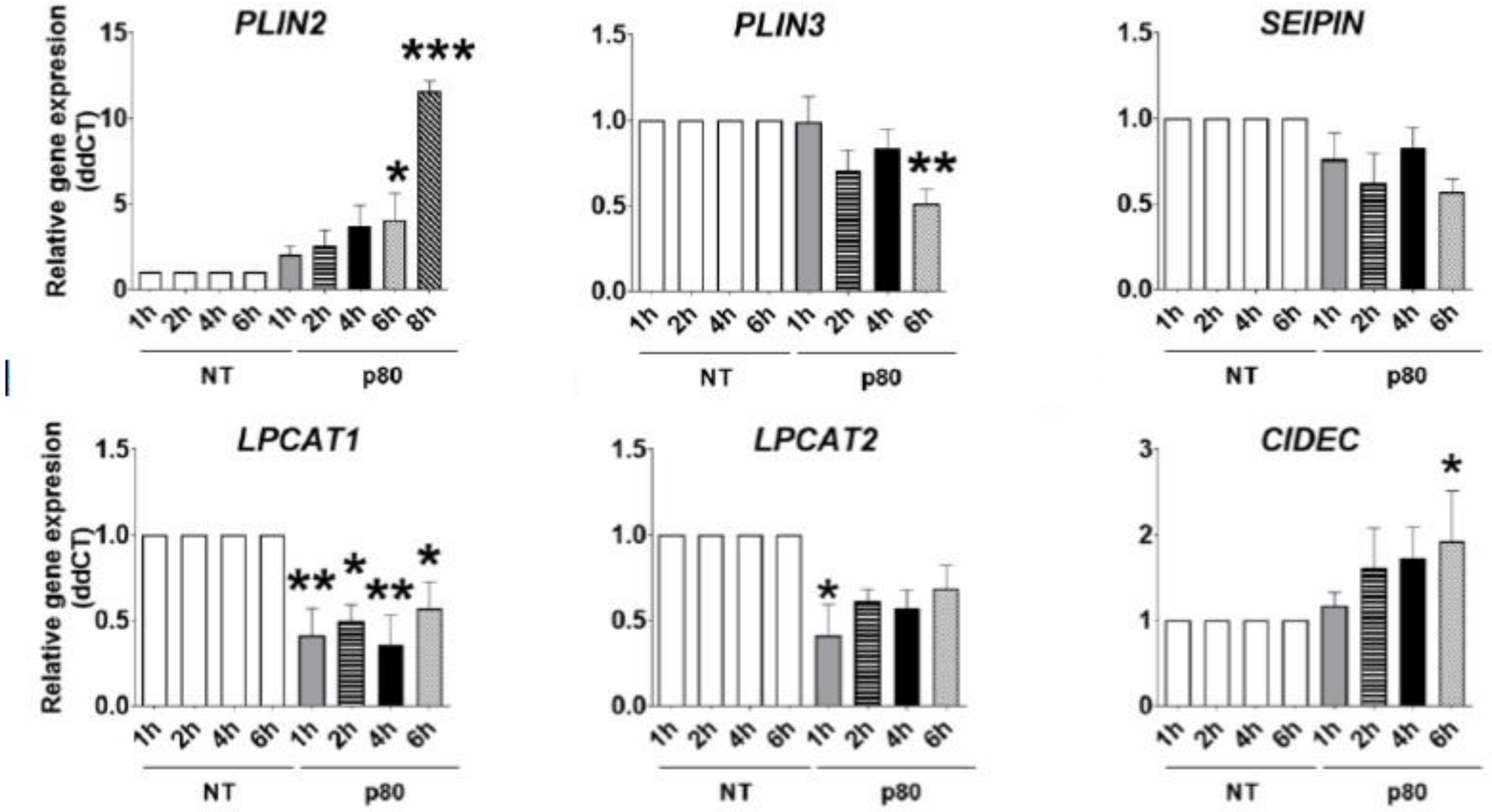

### Supplementary Figure 5

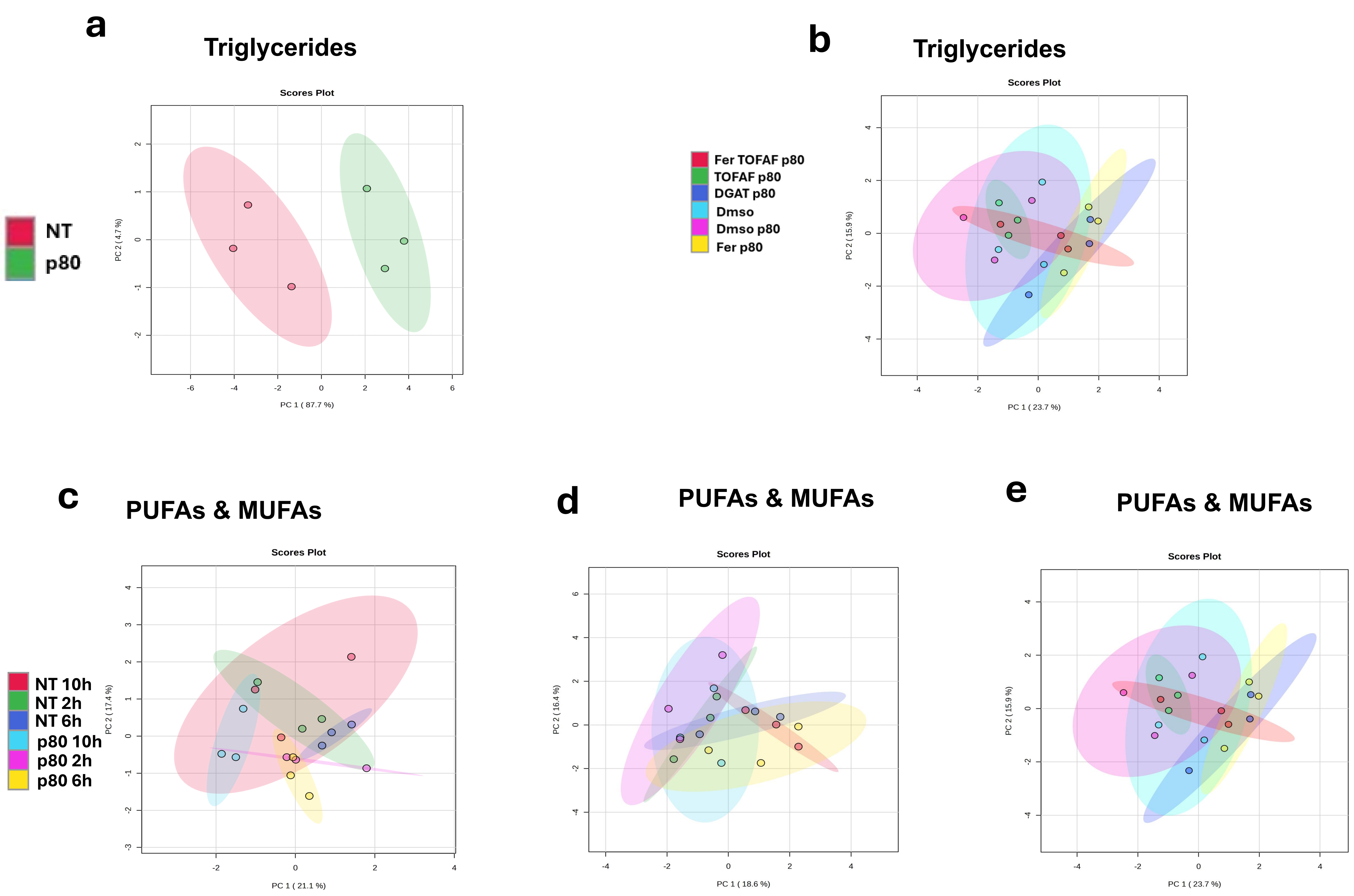

### Supplementary Figure 6

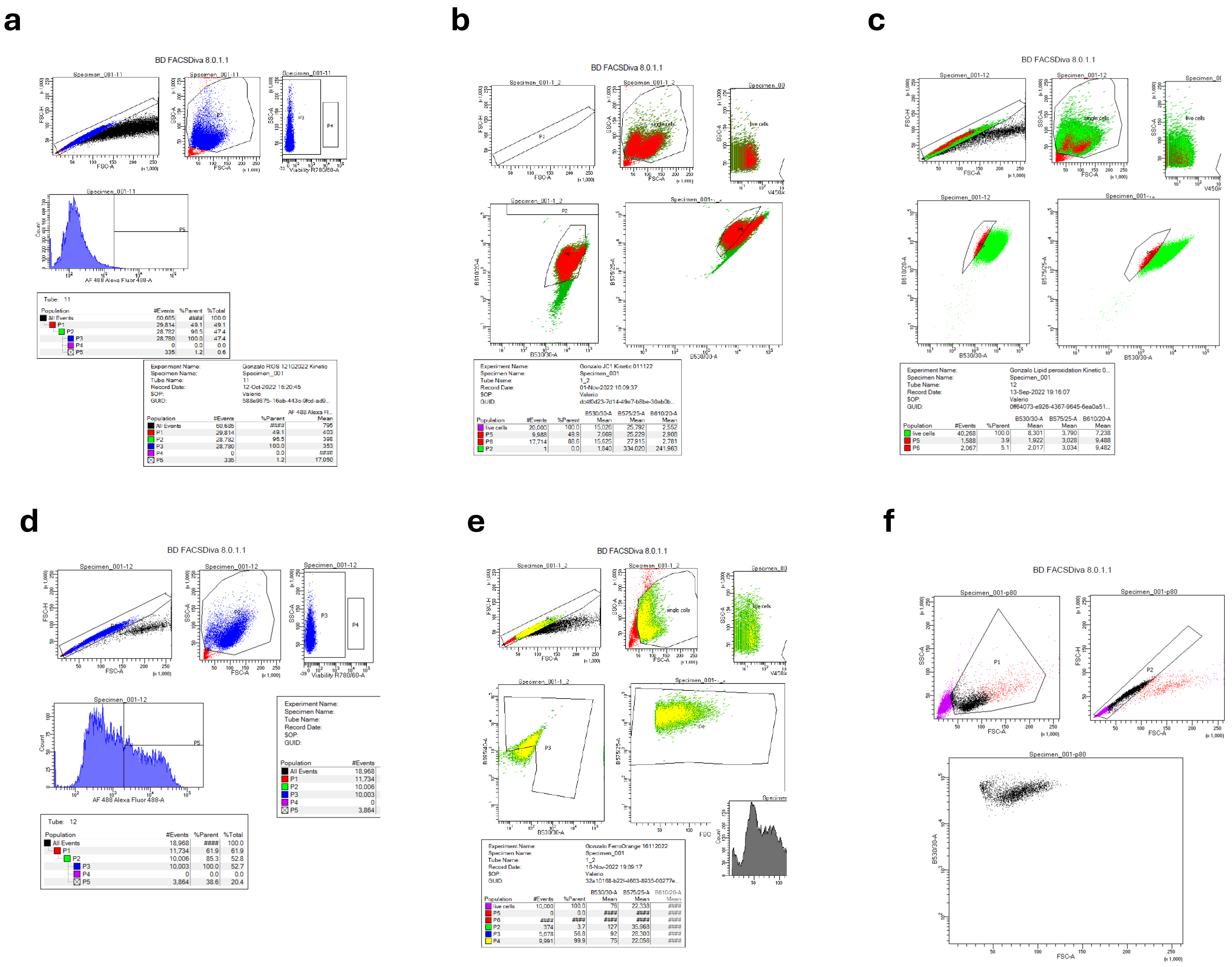

**Supplementary Figure 1. Cell viability and organoid growth after exposure to polysorbate-80 (p80) or carboxymethylcellulose (CMC).**

(a) HT29, C2BBe1, and DLD-1 cells treated with p80 for 6 h and 24 h; viability measured by CellTiter-Glo and expressed as % of time-matched NT (=100%). (b) HT29 cells treated with CMC (0.0625–0.5% w/v) for 1 h, 6 h, and 24 h; viability by CellTiter-Glo (top) and CellTiter-Blue (bottom), normalized to time-matched NT. (c) Bright-field micrographs of mouse small-intestinal organoids cultured ± 0.06% p80 over Days 2, 4, 6, and 8; scale bars, 400 µm. Bars show mean ± SEM of n = 3 independent experiments; statistics by one-way ANOVA with Dunnett's test vs NT (*p < 0.05; **p < 0.01; ***p < 0.001).

**Supplementary Figure 2. Induction of apoptotic markers and mitochondrial dysfunction by p80.**

(a) Caspase-3/7 activity at 1 h, 6 h, 10 h, and 24 h after 0.25% (v/v) p80; values are % of time-matched NT (=100%); mean ± SEM, n = 3 independent experiments. (b) qRT-PCR of apoptosis genes BAX, PUMA, and CASP3 after 24 h p80; expression normalized to housekeeping control and plotted relative to NT (=1); mean ± SEM, n = 3; two-tailed t-test vs NT. (c) Basal respiration and glycolysis prior to Mito Stress injections. Basal OCR (pmol $O_2 \cdot min^{-1}$) and ECAR ($mpH \cdot min^{-1}$) were taken from pre-injection cycles 1–3 (Agilent Wave) for NT and 0.25% p80. Bars also show non-mitochondrial OCR. Mean ± SEM (Plate 3: n = 3 wells/group; Plate 5: n = 2 wells/group); one-way ANOVA (Tukey). (d) JC-1 staining of mitochondrial membrane potential (ΔΨm): green = monomers, red = aggregates, nuclei Hoechst; representative fields at 1 h, 3 h, 6 h, and 10 h; scale bar, 50 µm.

**Supplementary Figure 3. ATP-based viability screen of compounds that mitigate p80 toxicity in HT29 cells.** (a) Rosiglitazone (PPARγ agonist). (b) Sulfasalazine (system $Xc^-$ inhibitor). (c) Fluvastatin (HMG-CoA reductase inhibitor). (d) Rosuvastatin (HMG-CoA reductase inhibitor). HT29 cells were co-treated with 0.25% (v/v) p80 and the indicated compound for 6 h or 24 h. Viability was measured by CellTiter-Glo and expressed as % of time-matched NT (=100%); DMSO is the vehicle control and "p80" denotes p80 alone. Bars show mean ± SEM (n = 3); statistics by one-way ANOVA (Dunnett) vs p80-only.

**Supplementary Figure 4. siRNA screen of lipid-handling genes in p80-treated HT29 cells.**

(a) Sytox Green membrane-permeability in cells transfected with siRNAs targeting (non-targeting siRNA = NT). Forty-eight hours post-transfection, cells were exposed to 0.25% (v/v) p80 for 1, 2, 3, 6, or 24 h with or without Ferrostatin-1 (Fer, 10 µM). Y-axis: Sytox signal (% of time-matched DMSO + NT-siRNA = 100%). Bars = mean ± SEM (n = 3); one-way ANOVA with Dunnett's test vs p80 + NT-siRNA, *p < 0.05, **p < 0.01, *** p < 0.001. (b) RT-qPCR of target genes PLIN2, PLIN3, SEIPIN (BSCL2), LPCAT1, LPCAT2, or CIDEC following 0.25% p80 treatment up to 6 h. Expression normalized to housekeeping gene *ACTB* (β-actin) and plotted relative to NT (=1) using the ΔΔCt method. Bars = mean ± SEM (n = 3); *p < 0.05, **p < 0.01.

**Supplementary Figure 5. PCA and triglyceride/FA remodeling after p80 in HT29 cells.**

(a) PCA scores plot of global lipidomic profiles separates NT and p80 (0.25% v/v, 24 h) samples; each point is a biological replicate. (b) Total triglycerides (TGs) after p80 vs NT; bars show median-normalized LC–MS peak area expressed as % of NT (=100%).

(c–e) Heatmaps of PUFA and MUFA species showing p80-induced changes (time points as indicated). Values are $\log_2$ fold-changes vs time-matched NT from aligned, median-normalized intensities; red = increase, blue = decrease (grey = not detected/not significant).

**Supplementary Figure 6. Flow cytometry gating and controls..**

Data were acquired on a BD FACSCelesta and analyzed in FACSDiva v8.0.1.1. For all assays, events were gated FSC-A/SSC-A (cells) → singlets (FSC-H vs FSC-A) → viable. Single-stain compensation and FMO + unstained controls were used to place gates. Plots are shown on bi-exponential scales; 1–2×$10^4$ viable events/sample. Gates and axes are kept identical across conditions for each assay; ratios are unitless and computed per cell (not dependent on total cell number).

(a) ROS ($H_2$DCFDA). Fluorescence collected on AF488-A (488-nm / 530/30). Panel shows the ROS-positive gate with NT vs 0.25% p80 overlays. Reported metric: % ROS-positive of viable singlets (mean ± SD). Percentages were analyzed after arcsine–square-root transform.

(b) Mitochondrial membrane potential (JC-1). Same upstream gating. JC-1 monomers (green) measured on AF488-A (530/30) and aggregates (red) on PE-TxRed-A (610/20). ΔΨm was quantified as per-cell red/green ratio = median(PE-TxRed 610/20 ÷ AF488 530/30) within the live-singlet gate; values were normalized to NT (=1). Representative red and green histograms/bivariates and gate positions are shown.

(c) Lipid peroxidation (BODIPY 581/591 C11). Same upstream gating. Oxidized C11 was detected in AF488 (530/30) and non-oxidized in PE (585/42). Reported metric: oxidized/non-oxidized ratio = median(AF488 ÷ PE) per cell, NT-normalized (=1). FMO and unstained controls are shown for gate placement.

(d) Mitochondrial $Fe^{2+}$ (Mito-FerroGreen). Same upstream gating. Signal recorded in AF488 (530/30). The Mito-FerroGreen-positive gate was set using AF488-FMO; representative NT vs p80 overlays are shown. Summary data are presented as % positive or AF488 MFI ($\log_{10}$-transformed) as indicated in the main figure.

(e) Cytosolic $Fe^{2+}$ (FerroOrange stain). Same upstream gating. Signal recorded in PE (B575/25-A; 488-nm excitation). The FerroOrange-positive gate was defined with PE-FMO and unstained controls. Representative NT vs 0.25% p80 overlays and % positive summary (mean ± SD) are shown; percent data were analyzed after arcsine–square-root transform.

(f) Neutral lipid droplets (BODIPY 493/503). Same upstream gating. Fluorescence collected in AF488 (530/30). Reported metric: % BODIPY-high cells of viable singlets; the “high” gate was positioned relative to NT using AF488-FMO. Where intensity is shown instead of % positive, values are MFI normalized to NT (=1).
